# Proteome-scale quantification of the interactions driving condensate formation of intrinsically disordered proteins

**DOI:** 10.1101/2024.12.21.629870

**Authors:** Rasmus K. Norrild, Sören von Bülow, Einar Halldórsson, Kresten Lindorff-Larsen, Joseph M. Rogers, Alexander K. Buell

## Abstract

Intrinsically disordered protein regions facilitate cellular organization through phase separation into biomolecular condensates. However, the molecular interactions driving this process remain poorly understood because of experimental limitations. Here, we advance experimental throughput by several orders of magnitude by developing Condensate Partitioning by mRNA-Display (CPmD). The method allows analysis of partitioning of hundred thousand peptides derived from the disordered proteome into reconstituted condensates. Our results demonstrate that the amino acid content, rather than specific sequence, primarily determines partitioning behavior. Importantly, quantification of the partitioning energies of peptides allows us to decipher the ‘molecular grammar’ of the relevant interactions, allowing accurate prediction of the formation of condensates of diverse full-length disordered protein regions. The results reveal how physicochemical properties of disordered regions encode biological functions through formation of biomolecular condensates.

## Introduction

Proteins are essential biomolecules that often require a well-defined three-dimensional structure to perform their functions. Therefore, the evolution of many protein genes is constrained by the need for a sufficiently large free energy difference between folded and unfolded states [1]. Precise structural organization in the folded state is required to be encoded in the amino acid sequence, which leads to conserved positional alignment of gene sequences from diverse species [2]. In contrast, Intrinsically Disordered Regions (IDRs) of proteins lack a well-defined folded state, which enables protein multifunctionality through diverse interaction networks and context-dependent structural adaptations essential for cellular regulation [3]. In addition to Short Linear Motifs (SLiMs) that can be aligned because they fold upon interaction, IDRs can also have a conserved composition and patterning of amino acids that are crucial for central biological functions [4, 5]. Recently, some IDRs have been shown to encode the localization and distribution of proteins to various subcellular compartments, including biomolecular condensates [6]. Exemplified by nucleoli, P-bodies, and germline granules, biomolecular condensates orchestrate cellular biochemistry and form through phase separation coupled to percolation of proteins and RNA [7–9]. Although multivalent motif-driven and stoichiometric interactions involving folded domains can lead to condensation, the polymeric and dynamic nature of IDRs makes them naturally prone to form condensates [10, 11].

NMR studies show that while all amino acids in the condensed phase have transient interactions, there are no dominant structures, raising questions about how biomolecular condensates form from IDRs [12–14]. In particular, it is unclear how multiple condensates coexist in the absence of structured interactions that have specificity [15, 16]. Transiently structured elements are present in most IDRs, and their role compared to that of amino acid composition and patterning in condensate formation remains controversial, in particular for transcription regulation and disease-related amyloid fibril formation [17–19].

Mutagenesis allows for the effective analysis of interactions driving condensate formation while circumventing the challenges of studying IDR dynamic structures with conventional structural biology methods [20, 21]. Such studies highlight the importance of aromatic amino acids and their distribution in the chain for condensate formation [22–25]. In addition, the presence of charged patches can contribute electrostatically to the driving force for condensate formation, in the absence of stably folded interactions [11, 26, 27]. However, the limited scalability of biophysical tests with purified IDR variants currently prevents a completely unbiased exploration of the ‘sequence grammar’ of IDR condensate formation [20, 21]. Therefore, conceptually new approaches are required for the study of IDRs at sufficient throughput.

In this study, we adapt protein display technology coupled to massively parallel DNA sequencing to study biomolecular condensate formation at an unprecedented scale. Although display technologies have generated high-quality data on protein folding and binding energetics for hundreds of thousands of sequences [28–30], they typically screen properties of individual molecules. However, condensate formation is a collective property requiring high concentrations of pure variants, which is a challenge for multiplexed assays. Inspired by a central idea of protein science, i.e. that partitioning of amino acids into nonaqueous solvents informs about their energetic contributions to protein folding, we developed Condensate Partitioning by mRNA-Display (CPmD). This approach measures peptide partitioning into reconstituted condensates as a scalable proxy for IDR condensate formation, as individual peptide fragments experience similar intermolecular interactions inside the condensates as they would within the context of longer IDRs [31]. We chose mRNA-display [32, 33] because both RNA and protein are natural components of biomolecular condensates and it is the minimally invasive peptide display technology: each peptide is covalently attached to only its encoding mRNA, via a flexible phosphate-containing polyethylene glycol (PEG) linker [34] (Methods, Supplementary Fig. 1).

We applied CPmD to the conserved RNA helicase DDX4, whose N-terminal IDR (DDX4N1) drives condensate formation [11, 15] to form essential germ granules that contain critical factors for germ cell determination [35–38]. Importantly, DDX4N1 remains disordered in both the dilute and condensed phases without converting to amyloid fibrils [12], making it an ideal model system for studying the molecular interactions governing condensate formation of IDRs.

## Results and discussion

### Peptide partitioning explains DDX4N1 condensate formation

We designed libraries with peptides of different lengths (14-50 aa) tiled through DDX4N1 starting at every second position, and one higher-resolution tiling using only peptides with 16 residues, but starting from every single position. Such tiling approaches have proven to be powerful in mapping important functional regions of IDRs [39]. We transcribed the libraries into mRNA constructs with and without displayed peptides, added the displayed and non-displayed mRNA libraries to the condensates in independent experiments, and initiated phase separation of reconstituted DDX4N1 condensates. By separating the dilute and condensed phases through centrifugation and sequencing the content of the dilute phase before and after induction of phase separation, we obtained partition free energies for all library members. We call this approach CPmD (Fig. 1a).

**Figure 1:**
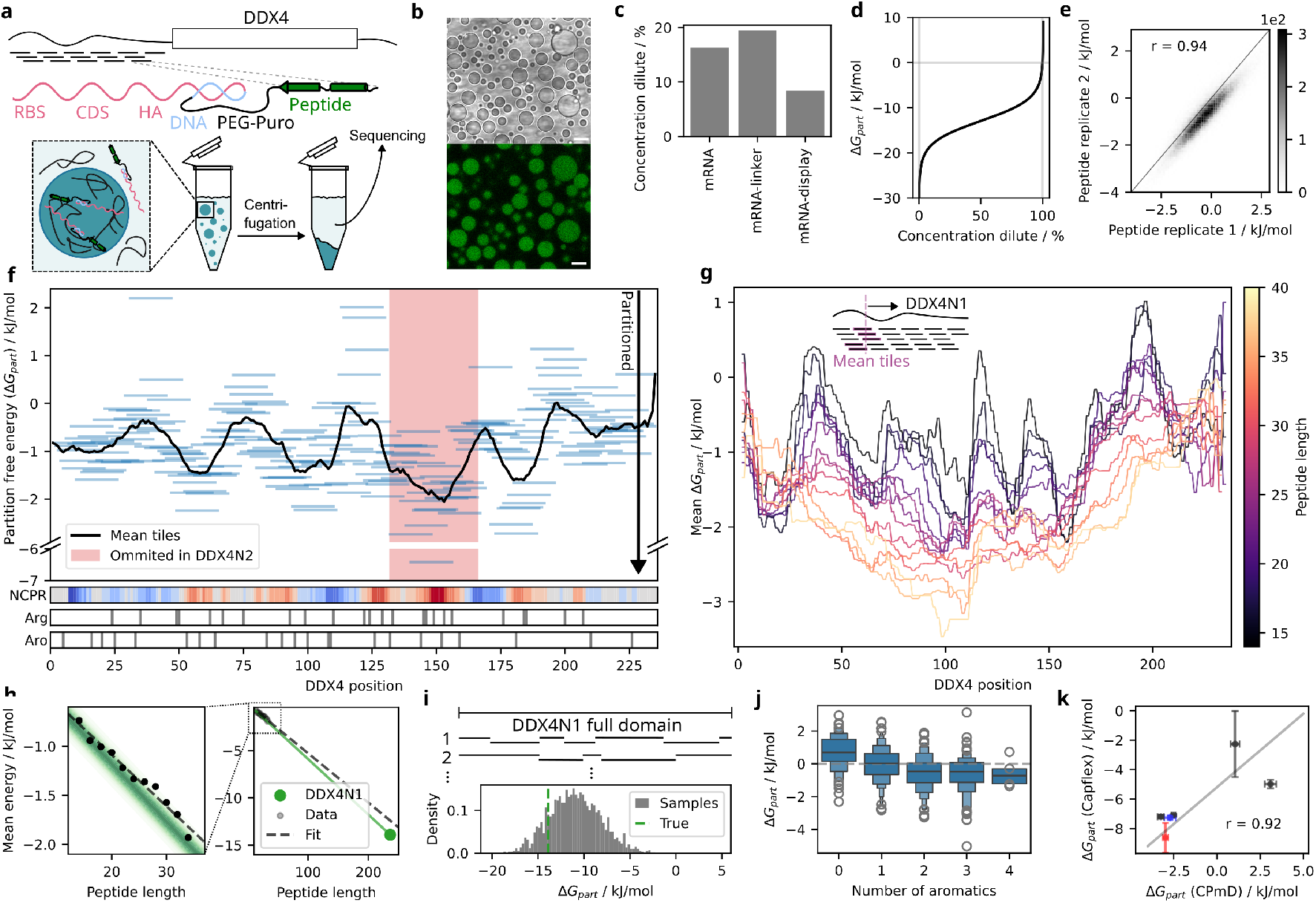
CPmD tiling of DDX4N1 quantitatively explains condensate formation. **a**, Workflow of Condensate Partitioning by mRNA-Display (CPmD) using peptides from DDX4’s N-terminal IDR. The mRNA (pink) is ligated to a puromycin-containing PEG-linker (black) with DNA sequence (blue) for hetero-duplex formation. **b**, Brightfield and fluorescence micrographs of DDX4N1 condensates with mRNA-display library containing a fluorescein-labeled puromycin linker. Scale bar: 10 *μ*m. **c**, qPCR quantification of mRNA-display library in the dilute phase after centrifugation. **d**, Partition free energy calculation using a dense phase volume fraction of 0.5%. **e**, Reproducibility of measured partition free energies between experimental repeats. **f**, Partition free energies of 16-amino-acid tiles showing a ∼40-amino-acid periodicity of interaction-prone regions. The DDX4N2 splice isoform lacking an exon (in red) containing the most partitioned region does not form condensates. **g**, Moving means of peptide tiles for different peptide lengths. **h**, Linear relationship between the mean partition free energy and peptide length, with expected dependence from full domain measurements (green). **i**, Partition free energy re-sampled as sums of different peptide tile sets. **j**, Strong correlation between the number of aromatic residues and peptide partitioning. **k**, Validation using synthetic peptides with Oregon Green fluorophore (OG488), showing agreement with mRNA-display measurements for negatively (blue) and positively (red) charged peptides.

The libraries were homogeneously distributed within the condensates, and even mRNA alone showed strong partitioning (Fig. 1b,c), consistent with previous findings that DDX4N1 condensates have a high affinity for single stranded nucleic acids [40]. Partitioning increased with higher purine content, lower secondary structure propensity, and longer sequences (Supplementary Fig. 3). Despite the strong mRNA partitioning, we isolated the free energy contributions to partitioning 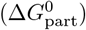 stemming from the peptides, by subtracting those from the RNAs and linker:

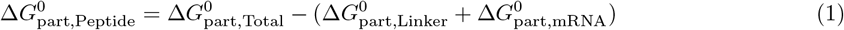

The partition free energies of the peptides were approximately 10% of the contribution of the mRNA (Equation 1, Fig. 1d,e), matching their relative molecular weights (Supplementary Fig. 1). The 16 amino acid DDX4N1 tiles partitioned with −1 to −2 kJ/mol free energy, representing only approximately 2-fold enrichment in the condensed phase (Fig. 1f). However, the length of the peptide and the average partition free energy showed a linear relationship in the area with reliable data (from 14 to 36 amino acids, Fig. 1g). Extrapolating to the full domain length (236 amino acids) yielded −12.7 kJ/mol, within error matching the expected energy based on measurements of the DDX4N1 domain’s dilute and dense phase concentrations alone (−13.9 kJ/mol) (Fig. 1h, Supplementary Fig. 4). Furthermore, we randomly selected peptide tiles of varying lengths that together make up the entire DDX4N1 domain. The sum of the partition free energies of these tiles successfully explained the condensate formation of the complete domain (Fig. 1i). These results confirmed that CPmD serves as a direct proxy for condensate formation.

### Specific regions of DDX4N1 drive condensation

A moving mean of the 16 amino acid tiles, where all partition energies of overlapping peptides are averaged along the sequence, revealed which regions of DDX4N1 drive condensate formation. The resulting profile showed a periodicity of ∼40 amino acids, indicating a block copolymer-like separation of regions driving partitioning and thus phase separation (Fig. 1f, Supplementary Fig. 4). Profiles derived from longer peptide tiles exhibited a stronger overall partitioning (Fig. 1g), consistent with the linear scaling of the partitioning. The resolution declined with increasing length, partly due to fewer sequencing reads (Supplementary Fig. 4). A central region of the IDR from position 140 to 160 demonstrated particularly strong partitioning. In support of CPmD’s relevance to condensate formation, the second natural isoform (UniProt ID: Q9NQI0-2), which lacks the exon encoding this region, does not form condensates under relevant conditions [9] (Supplementary Fig. 5). The center of the exon is strongly positively charged (Fig. 1f), and previous studies partly attributed condensate formation of DDX4N1 to electrostatic interactions between charged patches [11, 12, 26, 41]. Furthermore, given the predicted net charge of DDX4N1 of −3 at pH = 6.5, positively charged peptides were expected to partition more strongly into condensates. Indeed, predicted peptide charge showed a negative correlation with partition free energy (−0.13 kJ/mol per positive charge). Surprisingly, this correlation was driven primarily by arginine (−0.28 kJ/mol) rather than lysine (−0.04 kJ/mol, Supplementary Fig. 6), likely because the guanidine group of arginine has delocalized electrons and forms stacking interactions with other arginines and aromatic amino acids [42]. In line with this, the regions of the profile most strongly partitioned showed enrichment in aromatic amino acids, with an average effect of −0.43 kJ/mol per aromatic residue (Fig. 1j). Although highly negatively charged mRNA could potentially mask charge effects on partitioning, we found no discrepancy between CPmD and measured partition free energies of solid-state synthesized peptides using Capillary flow experiments (Capflex) [43] (Fig. 1k). Collectively, these findings demonstrate that CPmD provides a picture at unprecedented detail of how specific regions rich in aromatic residues and arginine drive intermolecular interactions in DDX4N1, explaining condensate formation primarily through hydrophobic interactions rather than electrostatic effects.

### Unbiased mutagenesis probes grammar of phase separation

We next asked what specific features of the protein sequence facilitated partitioning into its condensates. To determine whether interactions are driven by specific motifs [17], we measured the partitioning of peptides with scrambled amino acid sequences for each tile in the WT sequence (Fig. 2a). In general, these scrambled peptides partitioned similarly to the WT peptides, indicating that amino acid composition rather than specific motifs primarily determine partitioning. However, the central region of DDX4N1 showed differences between scrambled and WT peptides. We found no statistically significant correlation with previously reported features associated with condensate formation [44], such as helix propensity or the distribution of charged and hydrophobic residues. The only significant factor that influenced the impact of scrambling was the strength of the partitioning of the WT peptide, whether it was particularly strong or weak (Supplementary Fig. 6). To further explore this sequence dependence, we created a combinatorial mutant library of a 40-residues long peptide fragment (residues 100-139) in the center of DDX4N1. The scale of our CPmD experiment enabled unbiased mutagenesis, yielding reliable data on 27,269 peptides (20,165 without stop codon) containing an average of 8.1 substitutions per peptide (20% residues mutated) (Fig. 2b and Supplementary Fig. 7). Using Global Multi-Mutant Analysis (GMMA) [45], we disentangled the effects of all single substitutions in the peptide (represented as changes in partition free energy: ΔΔ*G*_part_) (Supplementary Fig. 8), revealing a resolvable degree of sequence-specific substitution effects (Fig. 2b). Nevertheless, the amino acid composition of mutated peptides remained the major determinant of their partitioning, demonstrated by strong correlation between experimental results and predictions based on the sum of amino acid partition free energies (Fig. 2c). When incorporating sequence position information, our predictions closely matched the experimental data within the limits of measurement noise. This shows that even in regions of DDX4N1 with strong sequence-specificity, approximately 80% of the partitioning behavior can be explained only by amino acid composition.

**Figure 2:**
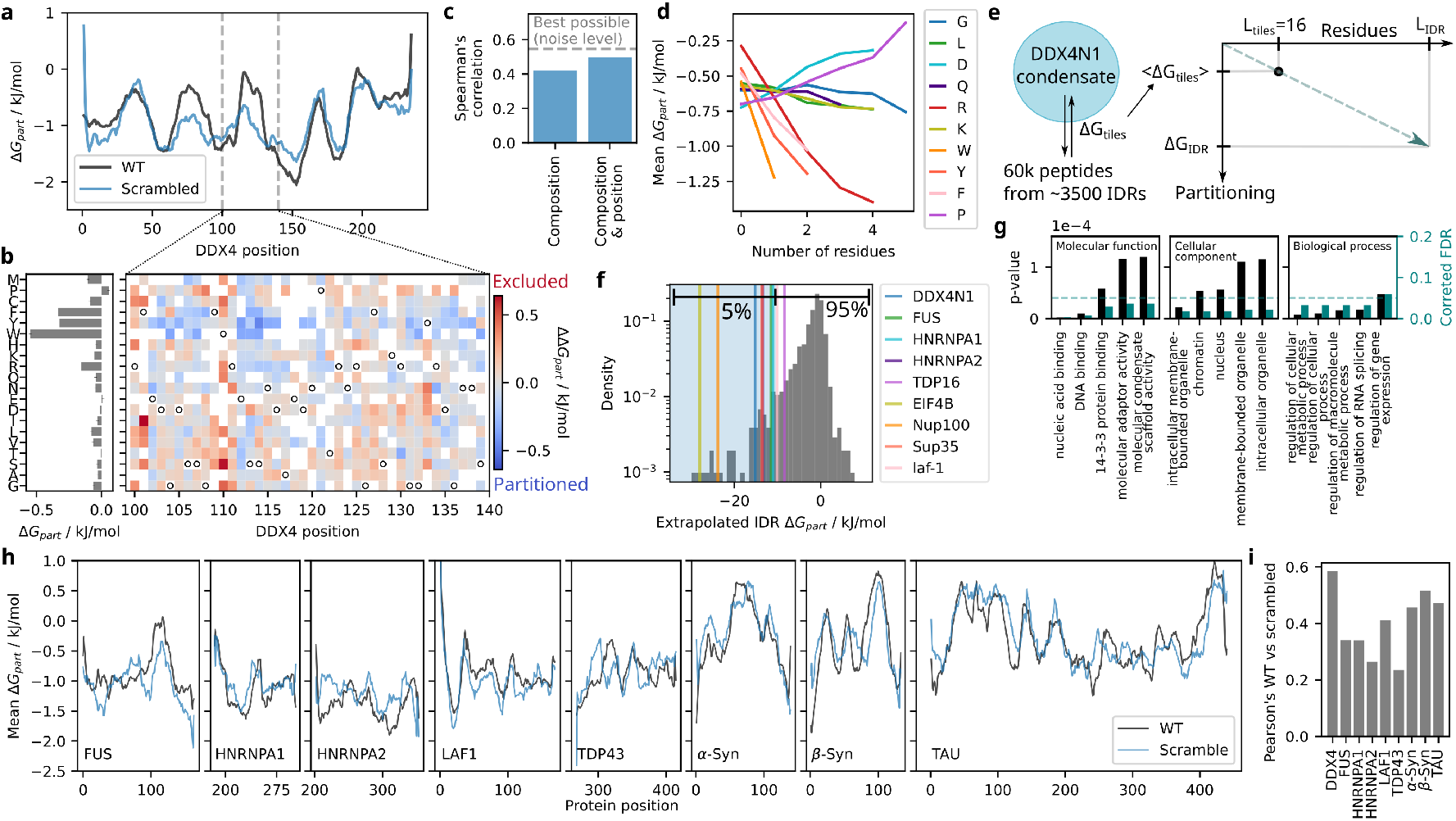
The same “sequence grammar” governs disordered proteome partitioning and DDX4N1 condensate formation. **a**, High-resolution tiling of 16-residue peptides of DDX4N1 into full-length condensates, with two scrambled peptides per tile to assess sequence-vs. composition-dependence. Moving means shown for both WT and scrambled peptides. **b**, Partition free energy changes (ΔΔ*G*_part_) measured across *>*20,000 multi-mutant variants of DDX4N1 residues 100-139. **c**, Comparison of linear models based on amino acid composition alone (left, 20 parameters) vs. composition and position (right, 800 parameters) for explaining multi-mutant data. **d**, Relationship between partition free energy and amino acid content in 16-residue peptides. **e**, CPmD measurement was done on peptide tiles across the entire DisProt database of experimentally verified IDRs. Full-length partition energies were obtained by extrapolation for all IDRs. **f**, Distribution of extrapolated partition free energies across the DisProt database, showing only 3% partition as strongly as DDX4N1 itself, and 7% would partition more than 50-fold. **g**, GO-term enrichment analysis of genes with strongly partitioning IDRs (*<* − 5.6 kJ/mol, 14% of data) reveals correlation with nucleic acid biology functions. **h**, Comparison of condensation propensity profiles between wild-type and scrambled sequences for known condensate-forming proteins. **i**, Correlation between partition free energies of wild-type peptides and their scrambled counterparts with identical amino acid composition.

### Other condensate forming IDRs partition to DDX4N1 condensates

In the absence of sequence motifs driving specific assembly, different chemical grammars have been proposed to govern the coexistence without mixing of diverse cellular condensates [15, 16]. We therefore investigated which IDRs of other proteins could partition into DDX4N1 condensates, and whether their amino acid composition differed from that governing DDX4N1 peptide partitioning. Capitalizing on the unique scale of CPmD, we measured a tiling library of 60,000 peptides that cover the disordered proteome of multiple organisms in the manually curated database of intrinsically disordered proteins (DisProt) [46].

Although this library partitioned significantly less than the DDX4N1 peptides on average, the dependence on amino acid composition remained similar, with increased hydrophobic amino acid content favoring partitioning (Fig. 2d, Supplementary Fig. 6). Building on the demonstrated linearity of partitioning with respect to length for DDX4N1 (Fig. 1h), we estimated partition free energies for all 3546 original full-length IDRs through extrapolation (Fig. 2e,f). The results showed that 15% of IDRs would be more than 10-fold enriched (Δ*G*_part_ *<* − 5.6 kJ/mol) in DDX4N1 condensates. This subset showed enrichment in gene ontology (GO) terms related to RNA-biology and biomolecular condensates (Fig. 2g). We observed only a weak correlation with proteins listed in the CD-CODE database of biomolecular condensates [47], primarily driven by proteins present in multiple condensates (Supplementary Fig. 6), consistent with recent observations [48]. This suggests that partitioning into DDX4N1 condensates relates either to the IDR being a condensate scaffold itself or to its recruitment to various condensates with low specificity.

37 human proteins in the library showed more than 25-fold enrichment in DDX4N1 condensates (Δ*G*_part_ *<* − 7.8 kJ/mol). 14 of these IDRs have been shown to form condensates in their purified forms and had diverse compositions (genes: AR, DDX3X, EIF4B, EWSR1, FUS, HNRNPA1, HNRNPA2B1, MAPT, POLR2A (RNA polymerase II C-terminal repeats) and TARDBP/TDP43 (literature references are listed in the Supplementary Information)). Another 11 IDRs were from proteins known to participate in condensate formation (literature references are listed in the Supplementary Information). The remaining 12 IDRs were from proteins that have no previously described condensate behavior (gene names: AHCTF1, AHR, CDKN2A, FGA, IBSP, LOX, NBN, NFATC2, RBM7, RICTOR, SECISBP2, and TP53BP2) (Fig. 2g). We also included a high-resolution sublibrary that contained 1-skip tiling, including scrambled peptides, of known condensate-forming proteins. These generally showed partitioning amplitudes of around –1 kJ/mol per 16 amino acid peptide, similar to DDX4N1 itself, indicating strong partitioning. However, these protein domains exhibited greater differences between WT and scrambled tiles compared to DDX4N1 (Fig. 2h,i), possibly due to transiently ordered segments influencing partitioning that are absent in DDX4N1. Notably, CPmD effectively captures this sequence specificity, enabling further study of how transient structure could encode condensate partitioning-specificity across diverse IDRs.

### CPmD reveals a core set of interactions driving condensate formation

Peptide fragments of DDX4N1 directly report on condensate formation of the full-length IDR, because they experience the same set of interactions in the condensate as the full-length sequence. However, peptides from other IDRs will experience different interactions in DDX4N1 condensates than during homotypic condensate formation. However, the partitioning of other condensate-forming IDRs to DDX4N1 condensates led us to investigate whether this behavior could provide general information on the formation of IDR condensates.

First, we examined this question using minimal perturbations caused by point mutations of DDX4N1 from our mutation library, introducing these mutations into purified full-length variants of the DDX4N1 domain. We quantified their homotypic phase separation using Taylor Dispersion-Induced Phase Separation (TDIPS) [49], which disperses salt ions and protein molecules differently in a microfluidic capillary, enabling robust and efficient exploration of phase separation behavior (Supplementary Fig. 5). Mutations showed stronger effects in TDIPS than in CPmD, indicating that homotypic condensate formation was more affected by mutations than partitioning into condensates formed by the WT sequence. We attributed this difference to the fact that substitutions simultaneously alter both the peptide’s intrinsic partition propensity and the host condensate’s general ability to recruit peptides.

In order to bridge the difference between the two scenarios, we developed a theoretical framework that disentangled these effects (Fig. 3a and Supplementary Fig. 9). We argued that the general ability of condensates to partition peptides should be related to the concentration of interaction-prone amino acids within them. A simple mean-field assumption, that a scaffold IDR’s propensity to recruit client peptides is directly proportional to its own energy density to partition to DDX4N1 (Δ*G*_Part_ per residue), reconciled differences between partitioning and intrinsic condensate formation (Fig. 3b and Supplementary Fig. 5).

**Figure 3:**
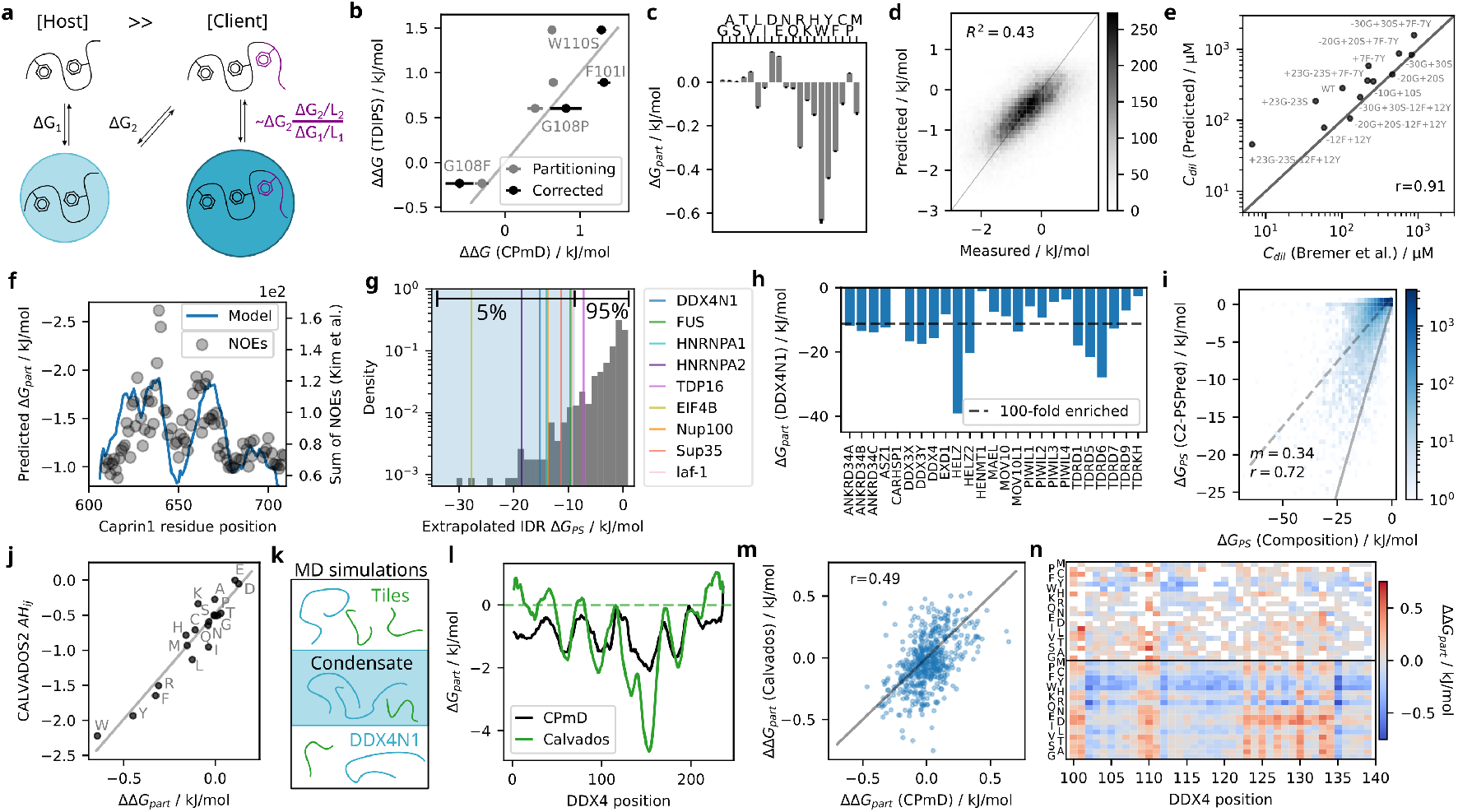
Peptide partitioning into DDX4N1 captures general properties of IDR condensate formation. **a**, Model for extrapolating intrinsic phase separation propensity from partitioning data, assuming condensate recruitment to be proportional to host peptide free energy density. **b**, Correlation between phase separation free energy changes measured by TDIPS and CPmD energies, with and without model correction. **c**, Amino acid transfer free energies fitted from DisProt library partitioning. **d**, Model performance evaluation. **e**, Composition-based model explaining dilute phase concentration differences in hnRNPA1-LCD mutations (20). **f**, Simulated tiling of CAPRIN1-LCD showing agreement with interaction hotspots identified by NOE signals (14). **g**, Estimation of intrinsic phase separation propensity for DisProt IDRs from CPmD tiling data after free energy density correction. **h**, Predicted partitioning of germ granule proteins into DDX4N1 condensates, with several showing *>*100-fold partitioning based on disordered regions. **i**, Comparison of human IDRome (50) phase separation predictions between composition model and PSPred. **j**, Correlation between CPmD-derived amino acid partition free energies and volume-integrated non-electrostatic potential energies in CALVADOS. **k**, CALVADOS simulations of CPmD experiments without mRNA and linker. **l**, Partition profiles comparison for DDX4N1 tiles from CPmD and simulations. **m**, Correlation of single-substitution effects between CPmD and simulations. **n**, Heatmap comparing single substitution effects in CPmD (top) and simulation data (bottom).

We then explored more broadly how the interactions that drive a peptide to partition into DDX4N1 condensates could represent the core set of interactions that govern IDR phase separation. We applied a similar approach to the composition model for the mutation library to analyze peptide partitioning across the DisProt database, yielding very similar amino acid specific parameters (Fig. 3c,d, Supplementary Fig. 6). With the mean-field correction described above, our model aligned remarkably well with published experimental data for mutation effects on phase separation of the unrelated hnRNPA1 low complexity domain at 20°C [20] (Pearson’s=0.91, Fig. 3e, Supplementary Methods) and predicted interaction hotspots in CAPRIN1-LCD that matched NMR measurements [14] (Fig. 3f).

Using this newly established quantitative link between partitioning into DDX4N1 condensates and *de novo* condensate formation, we converted our CPmD data on partitioning to represent homotypic condensate formation for all full-length IDRs in the DisProt library (Fig. 3g). Many of the strongly partitioned proteins mentioned above are also predicted to homotypically form condensates. Notably, these data accurately predict condensate formation of prion-like LCDs, such as FUS, hnRNPA1, hnRNPA2, TDP43, despite their amino acid composition differing significantly from that of DDX4N1. Our partitioning model also predicted partitioning to DDX4N1 condensates for IDRs absent from the DisProt library, revealing that several proteins expected to colocalize with DDX4N1 in germ granules contained IDRs predicted to cause strong partitioning (Fig. 3h).

To explore the generalizability of the composition model, we predicted the homotypic phase separation propensity of the entire human IDRome [50], and compared it with predictions from PSPred, a model trained from coarse-grained molecular dynamics simulations [44, 51] (Fig. 3i). We found a strong correlation with our composition-based model estimating a three-fold higher driving force for condensate formation compared to PSPred. PSPred is based on the CALVADOS force field, designed to simulate the structural ensembles of IDRs. The central parameters describing the “stickiness” of the coarse-grained amino acids (one bead per amino acid) were optimized using experimental data of IDR structural ensembles, with approximately 100 previously reported hydrophobicity scales serving as Bayesian priors [51]. We found that the amino acid-specific partition free energies of the composition model correlated extremely well with the “stickiness” parameters, shown as an integrated form of the pair potential AH_pairs_ [44] (Fig. 3j). This remarkable correlation underscores that the amino acid-specific transfer free energies in our partitioning model represent a fundamental physicochemical hydrophobicity scale, based on dynamic interactions between amino acids [52–54].

To test the agreement between CALVADOS simulations and our CPmD measurements, we simulated tiled peptide partitioning into excess full-length DDX4N1 (Fig. 3k and Supplementary Fig. 8). The individual peptides showed decent correlation (Pearson’s r = 0.63, Supplementary Fig. 8), which improved when taking the moving mean along DDX4N1 (Fig. 3l). We also simulated the partitioning of multimutant peptides of DDX4N1 that showed good correlation, especially on single amino acid-effects extracted using GMMA above (Fig. 3m,n, Supplementary Fig. 8), further validating our approach for deciphering the dynamic interaction network governing biomolecular condensate formation.

## Conclusions

In this study, we introduce Condensate Partitioning by mRNA-Display (CPmD), a high-throughput approach that provides unprecedented insight into the molecular grammar governing biomolecular condensate formation. Our proteome-scale measurements reveal that amino acid composition, rather than specific sequence motifs, primarily determines partitioning behavior in biomolecular condensates. This compositional code elucidates how intrinsically disordered regions, despite lacking well-defined structures, can encode complex biological functions through collective physicochemical properties.

The remarkable correlation between our experimentally determined amino acid partition free energies and the CALVADOS “stickiness” parameters underscores that our approach captures the dominant fundamental interactions governing IDR behavior. Inside a protein condensate, amino acids experience an averaged environment of all other amino acids, providing a more natural scenario than hydrophobic solvents used for traditional hydrophobicity scales. This explains why our partitioning experiments with a single condensate type yield insights with surprising generalizability across the disordered proteome.

The mean-field model we present extends beyond DDX4N1, quantitatively explaining phase separation of unrelated IDRs despite lacking data on peptide net charge, which was likely masked by the mRNA’s strong negative charge in CPmD. Protein net charge influences complex coacervation [55], and close-to-neutral proteins are generally less soluble, which also translates into a greater tendency to undergo homotypic phase separation [20, 51]. Although charge patterning can affect homotypic phase separation [11, 27, 44, 56], the charges in naturally occurring IDRs are generally well mixed [56, 57]. The predictive success of our charge-deficient mean-field model strongly supports the importance of generic hydrophobicity for biomolecular condensate formation. Furthermore, we find that substitution effects are largely additive for condensate formation, which parallels the principles observed in two-state protein folding [58, 59]. Analogously, our mean-field model thus elucidates a global constraint that might govern the evolution of the IDR sequence. The need to maintain the phase separation potential in the face of random mutations would retain a constant chemistry rather than sequence, which has been observed in IDRs [4, 16, 50].

As DNA synthesis and sequencing costs continue to decrease, CPmD will enable experiments of even greater scale and complexity. This approach opens possibilities for investigating reconstituted biomolecular condensates at near-cellular complexity, including folded protein domains. CPmD can also discover peptides capable of specific enrichment in condensates to enable detection or drug delivery. As condensates are increasingly recognized as central organizers of cellular biochemistry with implications in neurodegeneration, cancer, and viral infection, CPmD provides a quantitative foundation for understanding the sequence-to-function relationships that encode crucial biochemical roles of condensates in health and disease.

## Acknowledgements

We are grateful for the assistance with performing Illumina sequencing runs received from Marlene Danner Dalgaard from DTU Multi Assay Core and Malene Taarngaard Nørager and Anne Meldgaard Hansen from the Kennedy center at Rigshospitalet, Copenhagen. We also thank Soumik Ray for assisting with confocal microscopy. We thank Gerhard Steger for help with the RNA structure prediction. R.K.N., E.H., and A.K.B. would like to acknowledge the Villum Foundation (grant number: 35823), the Novo Nordisk Foundation (grant number: NNFSA170028392), and the European Union (ERC CoG 101088163 EMMA) for funding.

J.M.R. was supported by the Novo Nordisk Foundation (grant number: NNF19OC0054441). S.v.B. acknowledges support by the European Molecular Biology Organisation through Postdoctoral Fellowship grant ALTF 810-2022. K.L.-L. acknowledges support by the Novo Nordisk Foundation via the PRISM (Protein Interactions and Stability in Medicine and Genomics) centre (NNF18OC0033950). We acknowledge access to computational resources from the Biocomputing Core Facility at the Department of Biology, University of Copenhagen, from the Resource for Biomolecular Simulations (ROBUST; supported by the Novo Nordisk Foundation; NNF18OC0032608), and the Danish National Supercomputer for Life Sciences (Computerome).

## Author contributions

R.K.N. conceptualised, designed, performed and analysed most of the experiments, and wrote the manuscript.

S.v.B. did molecular dynamics simulations and analysis of these. E.H. performed and analysed experiments on purified recombinant proteins. K.L.-L. supervised and provided feedback at all stages of the project.

J.M.R. conceptualised and supervised mRNA-display experiments, and contributed to writing the manuscript.

A.K.B. conceived the project, supervised the work, contributed to writing the manuscript, and acquired funding. All authors commented on, edited, and approved of the manuscript.

## Competing interests

R.K.N., J.M.R., and A.K.B. share a part in a patent application based on the technology developed in this manuscript. K.L.-L. holds stock options in and is a consultant for Peptone. A.K.B. is a member of the scientific advisory board of Nuage Therapeutics.

## Data availability

Data and code are available on Zenodo at https://doi.org/10.5281/zenodo.14534152.

## Methods

### DDX4N1 CtoA expression and purification

A mutant of the N-terminal domain of Dead-box helicase 4 (DDX4N1) was used in which all cysteine residues were mutated to alanines (CtoA), as this simplifies handling of the protein and removes the potential for disulphide bridge formation and has little or no effect on LLPS[49]. Purification was performed through the recombinant expression of DDX4N1 CtoA fused to a *de novo*-designed thioredoxin-tag, eMM9[45]. Briefly, eMM9 is thought to facilitate protein expression through its high expression level and its high thermodynamic stability. The stability confers resistance to both heat and high denaturant concentrations useful for purification strategies used for intrinsically disordered proteins. A pET29(a) plasmid was constructed using Golden gate cloning containing an open reading frame coding for C-terminally His-tagged eMM9 followed by codon-optimized DDX4N1 CtoA linked by a 10-residue GS-linker and a TEV-cleavage site (ENLYFQ/G). The plasmid was used to transform *E. coli* strain BL21 (DE3) (New England Biolabs), and a clone was selected on kanamycin LB-agarose plates and Sanger-sequenced to confirm the correct sequence. 10 ml overnight culture was prepared in LB-medium supplied with 50 mg/l kanamycin and was used to inoculate 1l AB-LB (LB-medium with 5 g/l NaCl supplemented with 40 mM sodium/potassium phosphate (K/NaPi) buffer, pH 7.0, 15 mM ammonium sulphate, 50 mM NaCl, 2 mM MgCl2, 0.1 mM CaCl2, 3 nM FeCl3, 50 mg/l kanamycin). The culture was grown at 37°C to OD600 1.6 and induced with 1 mM IPTG before incubating overnight at 20°C. The next day, the cells were harvested by centrifugation at 7000 xg for 20 min at 4°C and the pellet was frozen and stored at -20°C. Next, 30 ml lysis buffer (50 mM NaPi buffer, pH 6.5 with 500 mM NaCl) was used to resuspend the cell pellet and 8 *μ*l Benzonase (purity *>* 90%, Merck-Millipore) were added. Cells were lysed by 30s/30s sonication pulses for a total time of 30 min on ice. Insoluble debris was then removed by centrifugation at 20000 xg for 20 min at 4°C and the supernatant was then heat-treated by incubation at 80°C for 15 min. Heat-induced protein aggregates were cleared by centrifugation at 4500 xg for 30 min at 4°C and passed through a 0.22 *μ*m filter. 1 M imidazole was added to a final concentration of 10 mM followed by quick mixing. The solution was then loaded onto 5 ml pre-equilibrated Ni-NTA resin (ThermoFisher Scientific) on a gravity flow column and washed with 10 column volumes of a buffer containing 50 mM NaPi, pH 6.5, 500 mM NaCl, and 20 mM imidazole, then 10 column volumes denaturing buffer containing 50 mM NaPi pH 6.5, 500 mM NaCl, 20 mM imidazole, and 3M GdnHCl. Finally, the denaturing salt was washed out with 10 column volumes of the non-denaturing wash buffer. A protease solution (9.5 ml lysis buffer with 2 mM TCEP and 500 *μ*l His-tagged TEV protease (10KU, Genscript or in-house purified)) was then added to release DDX4N1 CtoA into the solution during over-night incubation at room temperature with gentle mixing. TEV protease stays bound, and the eluted solution was concentrated using a 10 kDa centrifugal filter to 2 ml and SEC purified on a HiLoad Superdex 75 16/600 column equipped to an Ä KTA Pure system in a cooled cabinet at 8°C. Pooled fractions were concentrated to 500-600 *μ*M and aliquots were flash frozen in liquid nitrogen before storing at -80°C. The protein purity was checked at different stages of purification using SDS-PAGE and the protocol yielded ∼14 mg protein per litre culture.

### Plate based purification of mutated protein variants

Primers for generating mutants via Golden Gate cloning were ordered from IDT (Germany) at 100 *μ*M concentration TE buffer in a 96-well plate. The primers targeted 30 different substitutions and deletions of DDX4N1. They were then diluted to 1 *μ*M in MilliQ to use for PCR amplification of the pET29a plasmid encoding eMM9-tagged DDX4N1 CtoA. 20 *μ*L PCR reaction was set up using 1x Phusion master mix (F531L, ThermoFisher Scientific), 0.04 ng/*μ*L plasmid, and 50 pM primers. The PCR reaction was done in a Opentrons pipetting robot equipped with a thermal cyler (Opentrons, USA). 30 cycles were run with initial denaturation for 30 s at 98°C, final extension for 10 min at 72°C, and cycling through: 5 s at 98°C and 240 s at 72°C. The PCR reaction was assessed on 1% agar gel and the successfully produced PCR products were pooled together, 3.7 *μ*L of 27 products, totaling 100 *μ*L. The pooled PCR product was cleaned with QIAquick PCR clean-up kit(Qiagen, Netherlands). The product was then ligated via Golden Gate Assembly in an overnight ligation reaction at 37°C with the following final concentrations: 1000U T4 DNA ligase (M0202, NEB, USA), 1x T4 DNA ligase buffer, 30U Bsal-HFv2 (R3733, NEB, USA), 192 ng PCR product. The reaction was then inactivated by heating to 60°C for 5 min. 1 *μ*L of Dpn1 (FD1703, Thermo, USA) was added and incubated for 1 hour at 37°C, and then inactivated at 80°C for 5 min. 50 *μ*L of BL21 (DE3) cells (NEB, USA) were transformed with the product and colonies formed were transferred to a 96-well agar kanamycin plate to be sequenced by Eurofins Genomics (Germany) and to a deep well plate for glycerol stocks.

The sequencing resulted in 23 unique sequences of mutants that were then used for expression together with DDX4N1 CtoA as a control. This was done in two 24 deep-well plates (2×2 ml of each variant). 20 *μ*L of overnight cultures made from the glycerol stocks were transferred to 2 mL of Studier ZYM-5052 medium (Teknova, USA) with kanamycin. The cultures grew in a Eppendorf ThermoMixer shaking at 800 RPM at 25°C for 24 hours. Cells were harvested in 2 mL tubes by centrifugation at 7000 g for 10 min. The supernatant was discarded, and the 2 mL from the other plate was added and centrifuged again. 500 mL wash buffer (20 mM NaPi, 500 mM NaCl, 20 mM Imidazole, pH = 6.5) was added to resuspended cells. The lysis was then performed at 80°C for 20 min, followed by centrifugation for 20 min at 20,000 g to remove debris and aggregates. The supernatant was purified using 200 *μ*L NTA-beads (88222, Thermo, USA) in a filter plate using a vacuum manifold. After washing, 500 *μ*L of the supernatant was loaded onto the plate, allowed to sit for 10 min before vacuum was applied, and then washed 3x with 200 *μ*L wash buffer + 3M GdnHCl. Then, several more wash steps were performed with wash buffer without GdnHCl. Finally, 100 *μ*L of TEV cleavage solution (CombiTEVp 11 *μ*M (in-house purified) in wash buffer supplemented with 0.5 *μ*M TCEP) was loaded and incubated overnight. The protein was then eluted using three additional 100 *μ*L washes with wash buffer and concentrated with 30 kDa filters (UFC503096, Merck Millipore, USA) to 30 *μ*L. The produced proteins were quality checked using SDS-PAGE (Supplementary Fig. 5) and their hydrodynamic radius was measured using FIDA.

### TIDPS measurements of critical concentration

All measurements were performed on the Fida 1 instrument (Fida Biosystems, Denmark) using a 50 cm long capillary with a 75 *μ*m diameter for TDIPS, and the standard capillary of 1 m. Samples were injected from a compatible V-bottom 96-well plate and from a standard 50-vial holder, and detected using a UV detector with a 280 nm LED for excitation and a 300 nm long pass filter for emission. The signal detection was performed with 450 V detector gain with a recording frequency of 90 Hz.

**Table 1:** FIDA method used for measuring TDIPS curves of DDX4N1.

| Tray | Vial | Pressure (mbar) | Time (s) | Outlet | Measure | Comment |
| --- | --- | --- | --- | --- | --- | --- |
| 2 | 1 | 3500 | 45 | Variable | No | 1M NaOH Wash |
| 1 | Analyte | 3500 | 45 | Variable | No | Buffer wash |
| 1 | Indicator | 50 | 15 | Variable | No | Protein loading |
| 1 | Analyte | 300 | 90 | Variable | Yes | Mobilize and measure |

### Partition coefficients of solid-state synthesised peptides

To validate the mRNA-display approach, we bought six peptides with 14 residues containing an additional N-terminal cysteine residues with a maleamide Oregon Green 488 fluorophore (Schafer-N, Denmark). 100 *μ*M stock solutions were obtained by solubilising the peptides in MilliQ water, and were subsequently diluted to 100 nM for the experiment in a buffer containing 20 mM NaPi and 0.05%(V/V) Tween-20 pH 6.5. A 300 *μ*M stock of DDX4N1 was prepared by mixing 50:50 with a buffer containing 20 mM NaPi, 500 mM NaCl, 0.05%(V/V) Tween-20 pH 6.5. 16 *μ*L peptide stocks were thoroughly mixed with 4 *μ*L 300 *μ*M DDX4N1 for a final concentration of 80 nM peptide, 50 *μ*M DDX4N1, and 100 mM NaCl. The solution was immediately measured on the Fida 1 (Fida Biosystems, Denmark) at 20°C in a standard capillary (1 m and 75 *μ*m diameter) using the 480 nm LED detector with a gain of 420 V and a measuring frequency of 90 Hz. Control samples without protein were also measured and the change in the dilute phase was quantified. All samples were measured in duplicates. Using a condensed phase volume fraction of 0.22%, the partition free energy of the peptides was calculated. The quantification of spikes (signal from condensates) was in agreement with the data from the dilute phase concentration but was not used for direct quantification.

**Table 2:** FIDA method used for measuring Capflex of DDX4N1 with fluorescently labelled peptides.

| Tray | Vial | Pressure (mbar) | Time (s) | Outlet | Measure | Comment |
| --- | --- | --- | --- | --- | --- | --- |
| 2 | 1 | 3500 | 45 | Variable | No | 1M NaOH Wash |
| 2 | 2 | 3500 | 20 | Variable | No | MilliQ wash |
| 2 | 5 | 3500 | 30 | Variable | No | Buffer wash |
| 1 | Indicator | 300 | 300 | Variable | Yes | Measure sample |

### Microscopy

1 nM mRNA-display libraries containing a FITC dye were mixed with 120 *μ*M DDX4N1 at 100 mM NaCl in NaPi buffer pH 6.5 with 0.05% Tween-20. Brightfield and confocal micrographs where recorded using a Leica SP8 confocal microscope (Germany) using an 63x/1.40 oil immersion objective at the DTU bio-imaging core facility. Confocal images were obtained with a 16-bit depth. An excitation wavelength of 488 nm and emission channel of 500–600 nm was used. The gain was set to 1121V for the fluorescent channel and the sensitivity was set to 85% to detect the faint signal from the mRNA-display libraries. Images were recorded on rectangular glass coverslips 24×50 mm Menzel gläser which where plasma cleaned for 1 minute using the PDC-002-CE plasma cleaner from Harrick Plasma (USA). Plasma cleaned PDMS gaskets with 3 *μ*L circular wells were attached to the coverslips and used for imaging.

### mRNA-display

#### Fragment library from oligo pool

To investigate which parts of the DDX4N1 sequence participated most strongly in the condensation, we designed a tiling library consisting of many overlapping fragments of the protein domain. The library was designed with fragments that ranged from 14 amino acids in length and increased by two to 50 amino acids (l = 14,16,18… 50). For each length, the overlapping fragments were placed to start from every other amino acid, so that for 14 amino acid length fragments, each amino acid in DDX4N1 sufficiently far from the N and C-terminus was present in 7 different fragments. The library was synthesised by Twist Bioscience (USA) with 5’ and 3’ regions designed for PCR amplification. This PCR step is recommended by the supplier, but also allowed the attachment of the remaining 5’ and 3’ regions needed for mRNA-display[34]. The library was solubilised to 2.5 ng/*μ*L in TE buffer (10 mM Tris-HCl, 1 mM EDTA, pH=8) and 1 *μ*L was added for a total of 100 *μ*L PCR solution (0.5 *μ*M `Fivep_flank`, 0.5 *μ*M `Threep_flank_HA`, 1X Phusion HF buffer, 0.2 mM dNTPs, and 0.02 U Phusion (NEB M0530, USA)). 14 cycles were run with initial denaturation for 30 s at 98°C, final extension for 60 s at 72°C, and cycling through: 10 s at 98°C, 30 s at 56°C, and 40 s at 72°C.

#### Mutation library from doped oligo nucleotides

Doped oligos are oligo nucleotides synthesised with error, by adding a small amount of each of the other three nucleotides to each vial for synthesis. The resulting synthesised products will be highly diverse, with almost all molecules being unique, given a sufficiently high mutagenesis rate. We designed two such oligos, `Pep6_lib_fw` and `Pep6_lib_rev`, to generate a randomly mutated library of a central region in DDX4N1 positions [100:139]. The two oligos were designed with constant regions to anneal to `Fivep_flank` and `Threep_flank_HA` respectively for PCR amplification. Additionally, the two oligos overlapped with 15 base pairs at their 3’-ends allowing them to anneal and be extended to a double stranded product using the Klenow fragment polymerase (NEB M0210, USA). The extension reaction was performed in 100 *μ*L with 1 *μ*M of each oligo in NEBuffer 2 (50 mM NaCl, 10 mM Tris-HCl, 10 mM MgCl2, 1 mM DTT, pH 7.9). The oligos were annealed in the buffer but without dNTPs and polymerase by heating the solution to 94°C for 2 minutes and then ramping down the temperature at 0.2 °C/s down to 50°C. The solution was kept for 2 minutes and then ramped down to 37°C at the same speed. Then 2.5 *μ*L 10 mM dNTPs and 1 *μ*L Klenow fragment was added and the reaction was incubated at 37°C for 30 minutes. To limit the number of variants in the library, the 10 nM double-stranded product from the Klenow extension was diluted 10-fold (5 *μ*L in 45 *μ*L 0.05% triton X-100) 6 times to a theoretical maximum amount of molecules in the solution in 1 *μ*L of ∼600,000. A PCR reaction similar to the one described above for the tiling library was performed using GC buffer instead of HF buffer, which was run for 33 cycles.

#### Disprot library

A 60,000 member tiling library of the Disprot database (release 2022 12) containing 16 residue-long peptides with 4-fold overlap was ordered from Agilent (G7270A, USA). The library was solubilised in 40 *μ*L TE buffer yielding ∼1 *μ*M concentration. It was then diluted 1000-fold into a 100 *μ*L PCR solution as described above, and split in 2×50 *μ*L during PCR amplification. 13 cycles were run with initial denaturation for 30 s at 98°C, final extension for 60 s at 72°C, and cycling through: 5 s at 98°C, 15 s at 66°C, and 30 s at 72°C. The two samples were combined after the PCR reaction.

#### Transcription

Before transcription, the template DNA was cleaned using phenol-chloroform extraction. Briefly, 3M NaCl was added to the solution to a final concentration of 300 mM. For a 100 *μ*L solution, 100 *μ*L (same volume) phenol-cholorform-isoamyl alcohol (PCI) at a 25:24:1 ratio pH=6.5 was then added before vortexing and centrifuging at 10,000 g for 4 minutes. The top aqueous phase was then transferred to a new tube, and cleaned by adding 100 *μ*L chloroform-isoamyl alcohol (CI) at 24:1 ratio. Again, the solution was vortexed and centrifuged to separate the two phases. Finally, the aqueous phase was then transferred to a new tube and precipitated by adding ethanol to a final 70% (v/v). The pellet was washed with fresh 25 *μ*L of 70% ethanol, dried for a couple of minutes, and then re-solubilised in 10 *μ*L RNAse free water, for a theoretical maximum of 5 *μ*M (assuming complete PCR and no loss in PCI/CI extraction). Transcription was performed using the Ribomax kit (Promega, USA) in a 50 *μ*L reaction: 10 *μ*L 5X buffer, 15 *μ*L 100 mM rNTPs, 2.5 *μ*L DNA template, 0.5 *μ*L 1 mM DDT, 0.3 *μ*L RNAsin Plus (Promega, USA), 5 *μ*L Enzyme mix, and 16.7 *μ*L RNAse free water. The reaction was incubated at 37°C overnight (∼16 hours). To remove template DNA, 5.7 *μ*L DNAse buffer (0.4M Tris pH 8.0, 0.1M MgSO4, 0.01M CaCl2) and 1 *μ*L DNAse 1 (Thermo EN0525,

USA) were added and the solution was incubated for a maximum of 30 minutes at 37°C. To stop the reaction, EDTA pH=8 was added to a final concentration of 50 mM and NaCl was added to a final concentration of 300 mM. The RNA was then purified using PCI/CI as described above using a PCI solution with pH=4.5. Instead of precipitation, the aqueous phase after CI treatment was transferred to a NAP-5 column (Cytiva, USA) equilibrated with 10 mL RNAse-free water. If adding a 70 *μ*L sample, 430 *μ*L of RNAse-free water was then added. Finally, 500 *μ*L RNAse-free water was added to elute the mRNA. 0.3 *μ*L RNAsin was added to the tube before storing at -80°C.

#### Puromycin ligation

Ligation to the puromycin-containing linker was done in 50 *μ*L reaction containing 7.5 *μ*M `FPur-Linker` and 5 *μ*M mRNA. A 25 *μ*L solution with 2x concentration of the two (15 and 10 *μ*M, respectively) was heated to 95°C and slowly cooled to room temperature. Then, 5 *μ*L 10x T4 RNA ligase buffer and 5 *μ*L T4 ssRNA ligase 1 (NEB, USA) were added and the reaction was incubated for 1 hour at 37°C. 3 *μ*L of the reaction was loaded onto a PAGE gel containing TBE (Tris-Borate-EDTA), 8M Urea, 20%(v/v) formamide, and 8% acrylamide 19:1 to evaluate ligation. EDTA was added to the remainder of the reaction to a final concentration of 50 mM and NaCl to 300 mM, and the solution was purified using PCI/CI as described above using pH 6.5 PCI solution. For the Disprot library, additional measures were taken to purify the ligated product from unligated mRNA. 2 *μ*L 10% acetic acid was added to decrease the pH of the solution, together with 3 *μ*L formamide to a final 5% to inhibit RNA-RNA interactions. Using pH 6.5 PCI solution, ligated mRNA was found to be located between the lower phenol and upper aqueous phase. The aqueous phase was removed and the remainder was washed with 300 mM NaCl, which was then removed again after centrifugation. The purified product was released using 100 mM Tris-HCl pH 8.0 with 300 mM NaCl by vortexing. The aqueous phase was recovered after centrifugation and cleaned using CI as described above. The aqueous phase was then precipitated in 70% ethanol, washed with 70% ethanol, and solubilised in 7.5 *μ*L RNAse free water. This solution also contained un-ligated mRNA, however, this was removed in the HA-tag purification step.

#### Translation

Translation of the ligated mRNA was performed using the PureExpress ΔRF1,2,3 kit (NEB, USA). A 25 *μ*L reaction was performed using 10 *μ*L solution A, 7.5 *μ*L solution B, 5 *μ*L template, and 2.5 *μ*L RNAse free water. The solution was incubated for 1 hour at 37°C for translation to proceed. 3.25 *μ*L 100 mM Mg(OAc)2 pH=8 were added to the solution to improve the efficiency of the puromycin reaction during 30 min incubation at 37°C. Finally, 8.25 *μ*L 100 mM EDTA pH=8 were added to release the ribosomes during 30 minutes incubation at 37°C. The translation solution was then mixed 1:1 with 2X blocking buffer (100 mM HEPES-KOH pH 7.6, 400 mM NaCl, 0.1%(v/v) Tween-20, 0.1 g/L yeast RNA (Thermo, USA), 0.2 g/L acetylated BSA (Thermor, USA), and 5 *μ*M `GS3an_R36`) and cooled on ice. 40 *μ*L Pierce Anti-HA beads (Thermo, USA) was washed 3 times in ice cold wash buffer (50 mM HEPES-KOH pH 7.6, 200 mM NaCl, 0.05%(v/v) Tween-20, 1 *μ*M `GS3an_R36`) using magnetic separation. The blocked translation mixture was used to re-suspend the pelleted beads and incubated for 1 hour at 4°C with rotation to allow binding. The supernatant was then removed after magnetic separation and two rounds of washing were swiftly performed using ice-cold washing buffer. A final wash was performed with elution buffer (20 mM NaPi, 500 mM NaCl, 0.05%(v/v) Tween-20, and trace amounts of RNAsin (Promega, USA)), before eluting the bound molecules by adding 20 *μ*L elution buffer and incubating the beads at 80°C for 10 minutes. The solution was aliquoted and flash-frozen in liquid nitrogen. Quantification was performed using qPCR or fluorescence from the fluorescein on the puromycin linker yielding a typical concentration of ∼20 nM.

## Partitioning experiments

An aliquot of the library to be screened for partitioning was diluted 10-fold with low salt buffer (20 mM NaPi and 0.05%(v/v) Tween-20) such that this solution could induce coacervation of DDX4N1 CtoA. In parallel, the mRNA used to create the library was diluted to the same concentration, usually ∼2 nM, to be used as a control. Both of these solutions were cooled on ice and then added to two different PCR tubes, one containing 1.5 *μ*L 600 *μ*M DDX4N1 in elution buffer and one with 1.5 *μ*L elution buffer without protein, resulting in four tubes in total. All tubes were incubated for 1 minute at room temperature and then centrifuged for 10 minutes at 10,000 g in a pre-equilibrated centrifuge at 20°C. After centrifugation, 1 *μ*L of the supernatant from each tube was immediately added to the reverse transcription solution (1x SS-IV buffer, 0.25 mM dNTPs, 0.75 *μ*M `GS3an_R36`, 1.5 units of SS-IV reverse transcriptase (Thermo, USA), and trace amount of RNAsin (Promega,USA)). The reaction was proceeded at 50°C for 1 hour to ensure completion. The resulting solution was stored at -20°C.

## Sequencing

### Library preparation for sequencing

Attachment of Illumina adapter sequences was done through two consecutive PCR reactions. But first, the concentration of the template in each of the reverse transcription reactions was estimated using qPCR, such that over-cycling could be avoided. The first PCR reaction was performed in 50 *μ*L using 0.15 *μ*M of `Rd1T7g10M_F71` and `an13Rd2_R49`, 1X HF buffer, 0.2 mM dNTPs, 0.5 *μ*L reverse transcription solution, and 0.5 *μ*L Phusion polymerase (NEB, USA). A varying amount of cycles (usually 10-20) were run based on the number needed to get 0.05 *μ*M double-stranded product. The PCR program included an initial denaturation for 30 s at 98°C, final extension for 60 s at 72°C, and cycling through: 10 s at 98°C, 30 s at 52°C, and 40 s at 72°C. For NovaSeq analysis, phasing version of the two primers with added random nucleotides to frame-shift the reads were used, such that all PCR products had the same length, e.g. 2 extra bases at one end and 6 extra bases at the other for a total of 8. For the second reaction, 0.15 *μ*M of `Rd2N(701)P7_R52` and `P5S(502)Rd1_F57` were used in a similar PCR reaction using 2.5 *μ*L of the former PCR reaction as a template in a 50 *μ*L reaction. Each sample was assigned a unique combination of multiplexing barcodes in the two primers. Samples of FragLib2 were cleaned with QIAquick PCR clean-up kit (Qiagen, Netherlands) and sequenced using MiSeq v3 chip for 2×300 bps read at the DTU Multi Assay Core. Samples for the mutation library, Pep6Lib, were cleaned using Mag-Bind TotalPure NGS (Omega Bio-tek, USA) and sequenced using an SP Flow-cell on NovaSeq at the Kennedy center, Rigshospitalet Copenhagen.

### Processing of data

NGMerge[60] was used to merge paired-end reads using the default configuration. For the fragment library, all merged reads were mapped to variants expected in the library, allowing a single base pair discrepancy originating from sequencing errors, using a custom Python script. For the mutation library, the reads were cut to only contain the mutated area and unique sequences were then counted using custom code written in C++. As the library variation is similar to sequencing errors, the only way to remove sequencing noise is to require a minimum amount of reads. This was implemented by requiring all variants to have at least ten reads and be present in all samples.

### Analysis and modelling

#### Calculation of partition energy

The model rests on a number of assumptions which are expanded on in the supplementary materials but briefly mentioned here. 1) The relative abundance of sequencing reads is directly proportional to the concentration of each variant in the library. 2) The concentration in the condensate phase is exactly the molecules removed from the dilute phase divided by the volume fraction of the condensed phase. To avoid contamination between phases, we chose to measure only the concentrations of library members in the dilute phase (Fig. 1c) and infer the concentrations in the condensed phase using information on the total concentrations in the library and the dense phase volume fraction. 3) The system is at equilibrium, and the decrease in the concentration of a molecule in the dilute phase is directly due to its partition constant and, thus the free energy of partitioning. 4) The free energy of partitioning is a sum of the contribution from mRNA, linker, and peptide. Derivations of the formulae used to calculate the partition energy can be found in the Supplementary Information.

#### RNA structure prediction

RNAfold[61] was used to predict minimum free energy (MFE) secondary structures and their energies for all mRNAs in the tiling library using the default parameters. mRNA structures mimicking those expected with the puromycin linker ligated to the 3’-end of the mRNA were predicted by appending the linker DNA sequence to the RNA and changing the single thymine to uracil. For the mutation library, the structure of the mRNA for the unmutated “WT” species was calculated using default parameters and the “-p” option to also calculate the partition function.

### Fitting of linear models

Weighted ordinary least square regressions where used to explain partitioning of both peptides and RNA. Sequences were encoded either using their amino acid or nucleic acid composition in 20-member vectors or using one-hot encoding. These were all fitted in Python using linear algebra in Numpy. When needed, L2 regularisation (also called Tickhonov or Ridge regression) was used and regularisation strength was tuned using 5-fold cross validation. Potential over-fitting was evaluated on 10% held out data in all cases. Model performance and regularisation parameters are all reported in Supplementary Table 5.

### GO-term enrichment analysis

To obtain the extrapolated partition free energy of all IDRs in the Disprot library, the average partition energy of the tiling peptides within one IDR was calculated. This value was then devided by 16 (the number of amino acids in the peptides) and multiplied by the length of the IDR. To obtain a single value for proteins with multiple IDRs, the one with the highest affinity for the DDX4N1 condensates was choosen. GO-term enrichment was performed using GOATOOLS[62] in Python, using genes from species with more than 100 entries in the disprot database. TaxIDs: 9606 (Human), 10090 (Mouse), 10116 (Rat), 4932 (Yeast), 3702 (mouse-ear cress), 7227 (fruit fly), and 6239 (*Caenorhabditis elegans*). Enrichment p-values where calculated using Fishers exact t-test and correction for multiple testing was done using the Benjamini and Hochberg procedure (Corrected false discovery rate (FDR)).

### Coarse-grained molecular dynamic simulations

We performed coarse-grained molecular dynamics (MD) simulations with openMM 8.0[63] using the CALVA-DOS 2 force field, with force field parameters described elsewhere[51, 64]. Briefly described, the CALVADOS 2 force field represents IDRs using one bead per residue, with neighboring amino acid residues bonded via a harmonic potential. Salt-screened electrostatic interactions are represented via a Debye-Hückel potential. Non-electrostatic short-ranged interactions are described by a modified Lennard-Jones potential with residue type-specific sizes and stickiness values *λ*.

Each simulation was performed at constant molecule number, volume and temperature (Langevin integrator, T=293 K, friction coefficient=0.01 ps^−1^) at 175 mM ionic strength and pH 6.5 in a tetragonal, periodic simulation box with edge lengths 20 nm x 20 nm x 300 nm. For WT tiling simulations, WT fragment tiles of 16 residue length were generated along the DDX4N1 sequence, beginning from every residue (res. 1-16, 2-17, etc). We simulated a slab of 200 molecules of DDX4N1 together with one copy of all WT tiles in the same simulation box. The WT tiling simulations were run in triplicate. The partitioning free energy Δ*G*_part_ for a given fragment tile was calculated using the ratio of average number of molecules in the dilute phase of the simulation box and the average number of molecules in the DDX4N1 condensate (dense phase) across the trajectory. The boundaries between dense and dilute phase were determined from the DDX4N1 concentration profile as described previously[64]. Reported Δ*G*_part_ values are averages across the three independent simulation runs.

To estimate single substitution effects on fragment partitioning into DDX4N1, all possible single substitution variants of a peptide corresponding to DDX4N1 residues 100-139 were generated. Five copies of each of the 20 fragments corresponding to the WT and all single substitutions at a given residue position were simulated together with a slab of 200 DDX4N1 molecules. Each setup was simulated in four replicates, resulting in 4×40=160 simulations. Reported ΔΔ*G* values are calculated as the mean partitioning Δ*G* of all 5×4 copies of a given variant fragment minus the mean partitioning Δ*G* of all 5×4×20 control WT fragments in the simulations.

We simulated a subset of 200 of the multi-mutant fragments for DDX4N1 residues 100-139 that only contained a defined subset of substitutions in batches of 20 different fragments together with a slab of 200 DDX4N1 molecules, as above. GMMA analysis was performed on these simulated data using the linear models described above similarly to how they were used for CPmD data.

## Notes

### Summary of Updates

Central analysis has been updated and the manuscript has undergone substantial re-writing for conciseness and clarity.

## References

1. Bloom, J. D. et al. Thermodynamic prediction of protein neutrality. Proceedings of the National Academy of Sciences 102, 606–611 (2005).

2. Ahdritz, G. et al. OpenFold: Retraining AlphaFold2 yields new insights into its learning mechanisms and capacity for generalization en. preprint (Bioinformatics, Nov. 2022).

3. Wright, P. E. & Dyson, H. Intrinsically unstructured proteins: re-assessing the protein structure-function paradigm. en. Journal of Molecular Biology 293, 321–331 (Oct. 1999).

4. Zarin, T. et al. Proteome-wide signatures of function in highly diverged intrinsically disordered regions. en. eLife 8, e46883 (July 2019).

5. Boija, A. et al. Transcription Factors Activate Genes through the Phase-Separation Capacity of Their Activation Domains. en. Cell 175, 1842–1855.e16 (Dec. 2018).

6. Kilgore, H. R. et al. Protein codes promote selective subcellular compartmentalization. en. Science 387, 1095–1101 (Mar. 2025).

7. Mittag, T. & Pappu, R. V. A conceptual framework for understanding phase separation and addressing open questions and challenges. en. Molecular Cell 82, 2201–2214 (June 2022).

8. Banani, S. F., Lee, H. O., Hyman, A. A. & Rosen, M. K. Biomolecular condensates: organizers of cellular biochemistry. en. Nature Reviews Molecular Cell Biology 18, 285–298 (May 2017).

9. Shin, Y. & Brangwynne, C. P. Liquid phase condensation in cell physiology and disease. en. Science 357, eaaf4382 (Sept. 2017).

10. Li, P. et al. Phase transitions in the assembly of multivalent signalling proteins. en. Nature 483, 336–340 (Mar. 2012).

11. Nott, T. J. et al. Phase Transition of a Disordered Nuage Protein Generates Environmentally Responsive Membraneless Organelles. en. Molecular Cell 57, 936–947 (Mar. 2015).

12. Brady, J. P. et al. Structural and hydrodynamic properties of an intrinsically disordered region of a germ cell-specific protein on phase separation. en. Proceedings of the National Academy of Sciences 114, E8194–E8203 (Sept. 2017).

13. Murthy, A. C. et al. Molecular interactions contributing to FUS SYGQ LC-RGG phase separation and co-partitioning with RNA polymerase II heptads. en. Nature Structural & Molecular Biology 28, 923– 935 (Nov. 2021).

14. Kim, T. H. et al. Interaction hot spots for phase separation revealed by NMR studies of a CAPRIN1 condensed phase. en. Proceedings of the National Academy of Sciences 118, e2104897118 (June 2021).

15. Sabari, B. R., Hyman, A. A. & Hnisz, D. Functional specificity in biomolecular condensates revealed by genetic complementation. en. Nature Reviews Genetics (Oct. 2024).

16. Holehouse, A. S. & Alberti, S. Molecular determinants of condensate composition. en. Molecular Cell 85, 290–308 (Jan. 2025).

17. Hughes, M. P. et al. Atomic structures of low-complexity protein segments reveal kinked sheets that assemble networks. en. Science 359, 698–701 (Feb. 2018).

18. Garcia-Cabau, C. et al. Mis-splicing of a neuronal microexon promotes CPEB4 aggregation in ASD. en. Nature (Dec. 2024).

19. Bremer, A. et al. Reconciling competing models on the roles of condensates and soluble complexes in transcription factor function en. Nov. 2024.

20. Bremer, A. et al. Deciphering how naturally occurring sequence features impact the phase behaviours of disordered prion-like domains. en. Nature Chemistry 14, 196–207 (Feb. 2022).

21. Rekhi, S. et al. Expanding the molecular language of protein liquid–liquid phase separation. en. Nature Chemistry 16, 1113–1124 (July 2024).

22. Martin, E. W. et al. Valence and patterning of aromatic residues determine the phase behavior of prion-like domains. en. Science 367, 694–699 (Feb. 2020).

23. Franzmann, T. M. & Alberti, S. Prion-like low-complexity sequences: Key regulators of protein solubility and phase behavior. en. Journal of Biological Chemistry 294, 7128–7136 (May 2019).

24. Wang, J. et al. A Molecular Grammar Governing the Driving Forces for Phase Separation of Prion-like RNA Binding Proteins. en. Cell 174, 688–699.e16 (July 2018).

25. Quiroz, F. G. & Chilkoti, A. Sequence heuristics to encode phase behaviour in intrinsically disordered protein polymers. en. Nature Materials 14, 1164–1171 (Nov. 2015).

26. Paloni, M., Bailly, R., Ciandrini, L. & Barducci, A. Unraveling Molecular Interactions in Liquid–Liquid Phase Separation of Disordered Proteins by Atomistic Simulations. en. The Journal of Physical Chemistry B 124, 9009–9016 (Oct. 2020).

27. Pak, C. W. et al. Sequence Determinants of Intracellular Phase Separation by Complex Coacervation of a Disordered Protein. en. Molecular Cell 63, 72–85 (July 2016).

28. Olson, C. A., Wu, N. C. & Sun, R. A Comprehensive Biophysical Description of Pairwise Epistasis throughout an Entire Protein Domain. en. Current Biology 24, 2643–2651 (Nov. 2014).

29. Rogers, J. M., Passioura, T. & Suga, H. Nonproteinogenic deep mutational scanning of linear and cyclic peptides. en. Proceedings of the National Academy of Sciences 115, 10959–10964 (Oct. 2018).

30. Tsuboyama, K. et al. Mega-scale experimental analysis of protein folding stability in biology and design. en. Nature 620, 434–444 (Aug. 2023).

31. Ng, S. C. & Görlich, D. A simple thermodynamic description of phase separation of Nup98 FG domains. en. Nature Communications 13, 6172 (Oct. 2022).

32. Nemoto, N., Miyamoto-Sato, E., Husimi, Y. & Yanagawa, H. In vitro virus: Bonding of mRNA bearing puromycin at the 3-terminal end to the C-terminal end of its encoded protein on the ribosome in vitro. en. FEBS Letters 414, 405–408 (Sept. 1997).

33. Roberts, R. W. & Szostak, J. W. RNA-peptide fusions for the in vitro selection of peptides and proteins. en. Proceedings of the National Academy of Sciences 94, 12297–12302 (Nov. 1997).

34. Yamagishi, Y. et al. Natural Product-Like Macrocyclic N-Methyl-Peptide Inhibitors against a Ubiquitin Ligase Uncovered from a Ribosome-Expressed De Novo Library. Chemistry & Biology 18, 1562–1570 (Dec. 2011).

35. Brangwynne, C. P. et al. Germline P Granules Are Liquid Droplets That Localize by Controlled Dissolution/Condensation. en. Science 324, 1729–1732 (June 2009).

36. Tanaka, S. S. et al. The mouse homolog of Drosophila Vasa is required for the development of male germ cells. en. Genes & Development 14, 841–853 (Apr. 2000).

37. Dehghani, M. & Lasko, P. In vivo mapping of the functional regions of the DEAD-box helicase Vasa. en. Biology Open 4, 450–462 (Apr. 2015).

38. Yamazaki, H. et al. Bombyx Vasa sequesters transposon mRNAs in nuage via phase separation requiring RNA binding and self-association. en. Nature Communications 14, 1942 (Apr. 2023).

39. DelRosso, N. et al. Large-scale mapping and mutagenesis of human transcriptional effector domains. en. Nature (Apr. 2023).

40. Nott, T. J., Craggs, T. D. & Baldwin, A. J. Membraneless organelles can melt nucleic acid duplexes and act as biomolecular filters. en. Nature Chemistry 8, 569–575 (June 2016).

41. Crabtree, M. D. et al. Repulsive electrostatic interactions modulate dense and dilute phase properties of biomolecular condensates en. preprint (Cell Biology, Oct. 2020).

42. Vernon, R. M. et al. Pi-Pi contacts are an overlooked protein feature relevant to phase separation. en. eLife 7, e31486 (Feb. 2018).

43. Stender, E. G. P. et al. Capillary flow experiments for thermodynamic and kinetic characterization of protein liquid-liquid phase separation. en. Nature Communications 12, 7289 (Dec. 2021).

44. Von Bülow, S., Tesei, G., Zaidi, F. K., Mittag, T. & Lindorff-Larsen, K. Prediction of phase-separation propensities of disordered proteins from sequence. en. Proceedings of the National Academy of Sciences 122, e2417920122 (Apr. 2025).

45. Norrild, R. K. et al. Increasing protein stability by inferring substitution effects from high-throughput experiments. en. Cell Reports Methods, 100333 (Nov. 2022).

46. Quaglia, F. et al. DisProt in 2022: improved quality and accessibility of protein intrinsic disorder annotation. en. Nucleic Acids Research 50, D480–D487 (Jan. 2022).

47. Rostam, N. et al. CD-CODE: crowdsourcing condensate database and encyclopedia. en. Nature Methods 20, 673–676 (May 2023).

48. Hadarovich, A. et al. PICNIC accurately predicts condensate-forming proteins regardless of their structural disorder across organisms. en. Nature Communications 15, 10668 (Dec. 2024).

49. Norrild, R. K. et al. Taylor Dispersion-Induced Phase Separation for the Efficient Characterisation of Protein Condensate Formation. en. Angewandte Chemie International Edition 63, e202404018 (June 2024).

50. Tesei, G. et al. Conformational ensembles of the human intrinsically disordered proteome. en. Nature 626, 897–904 (Feb. 2024).

51. Tesei, G., Schulze, T. K., Crehuet, R. & Lindorff-Larsen, K. Accurate model of liquid–liquid phase behavior of intrinsically disordered proteins from optimization of single-chain properties. en. Proceedings of the National Academy of Sciences 118, e2111696118 (Nov. 2021).

52. Rekhi, S. & Mittal, J. Amino Acid Transfer Free Energies Reveal Thermodynamic Driving Forces in Biomolecular Condensate Formation en. Dec. 2024.

53. Ambadi Thody, S. et al. Small-molecule properties define partitioning into biomolecular condensates. en. Nature Chemistry (Sept. 2024).

54. Urry, D. W. et al. Temperature of polypeptide inverse temperature transition depends on mean residue hydrophobicity. en. Journal of the American Chemical Society 113, 4346–4348 (May 1991).

55. Aumiller, W. M. & Keating, C. D. Phosphorylation-mediated RNA/peptide complex coacervation as a model for intracellular liquid organelles. en. Nature Chemistry 8, 129–137 (Feb. 2016).

56. Pesce, F. et al. Design of intrinsically disordered protein variants with diverse structural properties. en. Science Advances 10, eadm9926 (Aug. 2024).

57. Holehouse, A. S., Das, R. K., Ahad, J. N., Richardson, M. O. & Pappu, R. V. CIDER: Resources to Analyze Sequence-Ensemble Relationships of Intrinsically Disordered Proteins. en. Biophysical Journal 112, 16–21 (Jan. 2017).

58. Wells, J. A. Additivity of mutational effects in proteins. en. Biochemistry 29, 8509–8517 (Sept. 1990).

59. Faure, A. J. et al. The genetic architecture of protein stability. en. Nature (Sept. 2024).

60. Gaspar, J. M. NGmerge: merging paired-end reads via novel empirically-derived models of sequencing errors. en. BMC Bioinformatics 19, 536 (Dec. 2018).

61. Mathews, D. H. et al. Incorporating chemical modification constraints into a dynamic programming algorithm for prediction of RNA secondary structure. en. Proceedings of the National Academy of Sciences 101, 7287–7292 (May 2004).

62. Klopfenstein, D. V. et al. GOATOOLS: A Python library for Gene Ontology analyses. en. Scientific Reports 8, 10872 (July 2018).

63. Eastman, P. et al. OpenMM 7: Rapid development of high performance algorithms for molecular dynamics. en. PLOS Computational Biology 13 (ed Gentleman, R.) e1005659 (July 2017).

64. Tesei, G. & Lindorff-Larsen, K. Improved predictions of phase behaviour of intrinsically disordered proteins by tuning the interaction range. en. Open Research Europe 2, 94 (Jan. 2023).

