## Supplementary Information for "Proteome-scale quantification of the interactions driving condensate formation of intrinsically disordered proteins"

Rasmus K. Norrild[1], Sören von Bülow[2], Einar Halldórsson[1], Kresten Lindorff-Larsen[2], Joseph M. Rogers[3,*], and Alexander K. Buell[1,*]

[1]Department of Biotechnology and Biomedicine, Technical University of Denmark, Kgs. Lyngby, Denmark
[2]Structural Biology and NMR Laboratory & the Linderstrøm-Lang Centre for Protein Science, Department of Biology, University of Copenhagen, Copenhagen, Denmark
[3]Department of Drug Design and Pharmacology, University of Copenhagen, Copenhagen, Denmark

May 2, 2025

#### Materials

Puromycin linker (`/5Phos/CTCCCGCCCCCCG/iFluorT/CC/iSp18//iSp18//iSp18//iSp18//iSp18/CC/3Puro/`) called `FPur-linker` was ordered from IDT (Germany). DNA primers were ordered from TAG Copenhagen (Denmark). Double distilled water was used for all RNA work (Invitrogen UltraPure DNase/RNase-Free Distilled Water).

$NaH_2PO_4$, $Na_2HPO_4$, Immidazole, $FeCl_3{\cdot}6H_2O$, $CaCl_2{\cdot}2H_2O$, $MgCl_2{\cdot}6H_2O$, $K_2HPO_4$, and $KH_2PO_4$ were all perchased from Sigma (USA). LB media and $(NH4)_2SO_4$ were purchased from VWR (USA). Kanamycin sulfate and NaCl was purchased from PanReac AppliChem (Germany).

#### Software

Mainly open-source and free software has been used, and we are very thankful to the community of people who create and maintain this powerful suite of tools. Select programmes and computer coding languages used are; Latex, Python, Inkscape, Gimp, C++, and Ubuntu/Linux. Data analysis was performed with Python version 3.11.7 using the following packages: Numpy v1.26.4, Pandas v2.2.3, Matplotlib v3.8.4, Seaborn v0.13.2, Scipy v1.14.0, Scikit-learn v1.5.1, Biopython v1.83, Uncertainties v3.2.2, Emcee v3.1.5, Metapredict v2.63[1], and localCIDER v0.1.21[2].

### Supplementary Text

#### Calculating partition energies from sequencing reads

Using the assumptions stated in the methods section of the main manuscript, we can first calculate the concentration of each library variant, $v$, in both the control and in the dilute phase after phase separation, $[v]_X$, by calculating the fraction of reads $F_{v,X}$. The different samples or sequencing runs are indicated by $X$:

$$F_{v,X} = \frac{\text{reads}_{v,X}}{\sum_i^N \text{reads}_{i,X}} \tag{S1}$$

The concentration of each member of the library is then calculated using the total concentration of the library $c_{\text{library},X}$ measured by qPCR directly during the experiment:

$$[v]_X = F_{v,X} \cdot c_{\text{qPCR},X} \tag{S2}$$

The fractional concentration in the dilute phase is then (using the short notation that the fractional concentration in the dilute phase of the whole library is called $f_{\text{qPCR}} = \frac{c_{\text{qPCR,dilute}}}{c_{\text{qPCR,total}}}$:

$$f_{D,v} = \frac{[v]_{\text{dilute}}}{[v]_{\text{total}}} = \frac{F_{v,\text{dilute}} \cdot c_{\text{qPCR,dilute}}}{F_{v,\text{total}} \cdot c_{\text{qPCR,total}}} = \frac{F_{v,\text{dilute}}}{F_{v,\text{total}}} f_{\text{qPCR}} \tag{S3}$$

To find the fractional total concentration in the condensed phase $f_{C,v} = \frac{[v]_{\text{condensed}}}{[v]_{\text{total}}}$, the overall conservation of amount of substance is used:

$$n_{tot} = n_C + n_D \tag{S4}$$

Using $n = Vc$, and collecting the concentration in the condensed phase on one side of the equation, $c_C$, we obtain

$$c_C = \frac{c_{\text{tot}} V_{\text{tot}}}{V_C} - c_D \frac{V_D}{V_C} \tag{S5}$$

Re-writing and introducing the fractional concentrations $f_x = \frac{c_x}{c_{\text{tot}}}$ yields:

$$c_{\text{tot}} f_C = c_{\text{tot}} + c_{\text{tot}} \frac{V_D}{V_C} - f_D c_{tot} \frac{V_D}{V_C} \tag{S6}$$

$c_{\text{tot}}$ then cancels out, and by re-writing and using fractional volume $\psi_C = \frac{V_C}{V_C + V_D}$ instead the equation can be written as:

$$f_C = \frac{\psi_C f_D + f_D + 1}{\psi_C} \tag{S7}$$

As equilibrium is reached when the chemical potential of the variant, $v$, is equal in both phases, the partition constant of each variant is thus the ratio between its activity in both phases. Here, we use the convention of 1 M standard state which is omitted for clarity:

$$(K_D^\circ)_v = \frac{a_{v,\text{condensed}}}{a_{v,\text{dilute}}} = \frac{[v]_{\text{condensed}}}{[v]_{\text{dilute}}} = \frac{f_C [v]_{\text{total}}}{f_D [v]_{\text{total}}} = \frac{\frac{\psi_C f_D - f_D + 1}{\psi_C}}{f_D} = 1 - \frac{f_D - 1}{\psi_C f_D} \tag{S8}$$

At constant pressure, we can thus calculate the standard Gibbs free energy of transfer:

$$\Delta G_v^\circ = -RTln(K_D^\circ)_v \tag{S9}$$

We then assume that the total partitioning free energy is a sum of the individual parts of the molecule: mRNA, linker, and peptide.

$$\Delta G^\circ_{\text{total}} = \Delta G^\circ_{\text{mRNA}} + \Delta G^\circ_{\text{linker}} + \Delta G^\circ_{\text{peptide}} \tag{S10}$$

For the first two libraries (FragLib and MutantLib) we were not yet able to isolate the mRNA ligated to the linker with sufficient purity. We were only able to measure the partitioning of untranslated mRNA alone. All reported effects of the peptide are therefore offset with a constant contribution from the linker.

$$\Delta G^\circ_{\text{app-peptide}} = \Delta G^\circ_{\text{total}} - \Delta G^\circ_{\text{mRNA}} = \Delta G^\circ_{\text{linker}} + \Delta G^\circ_{\text{peptide}} \tag{S11}$$

Note that normalisation by mRNA also has the effect that the contribution from the fractional volume of condensed phase is effectively eliminated from the calculated energy if both $f_D$ and $\psi_C$ are small because $K_D^\circ \approx -\frac{f_D - 1}{\psi_C f_D} = (\frac{1}{f_D} - 1)\frac{1}{\psi_C}$ (see equation S8) and therefore cancels out:

$$\Delta G_1^\circ - \Delta G_2^\circ \propto ln(K_D^\circ)_1 - ln(K_D^\circ)_2 = ln\frac{(K_D^\circ)_1}{(K_D^\circ)_2} = ln\frac{(\frac{1}{f_{D,1}} - 1)\frac{1}{\psi_C}}{(\frac{1}{f_{D,2}} - 1)\frac{1}{\psi_C}1} = ln\frac{\frac{1}{f_{D,1}} - 1}{\frac{1}{f_{D,2}} - 1} \quad \text{(S12)}$$

For the DisProt library, the obtained library concentrations, $c_{\mathrm{qPCR}}$, had to be corrected, because the resulting concentrations of a number of library members, $[x]_X$, were physically impossible. Several other partition free energies were highly skewed because they ended up in the tails of the logarithmic function in equation S9. The qPCR measurements are expected to have quite high errors (in logarithmic space because of the nature of the PCR reaction), which necessitated us to make corrections to the values. We therefore devised a strategy to correct these values by the minimum amount necessary, such that no peptides in the library had impossible concentrations. This likely leaves some data in the extreme regions of S9, but further corrections would be highly subjective.

Error estimates were constructed based on Poisson errors, $\sigma(\mathrm{reads}_{v,X}) = \sqrt{\mathrm{reads}_{v,X}}$, and propagated through to the final estimates of partitioning free energy using the `uncertainties` module in Python.

### Estimation of condensed phase volume fraction

We relied on a phase diagram determined by Brady *et al.*[3] to estimate the volume of the condensed phase. The data was fitted to an approximate analytical solution of the Flory-Huggins model for polymer phase separation, using a density of 1350 g/L for a pure protein [4]. Using the dilute phase concentrations measured for Ddx4N1 at 140 and 158 mM NaCl, 36 and 46 $\mu$M respectively, the estimated concentrations in the condensed phase were 17.2 and 16.8 mM. This resulted in the dense phase volume fractions being 0.49% and 0.44%, respectively. As described above, these values mostly canceled out when calculating differences in partition energies.

##### Linear model of amino acid effects

Three different models were used to capture peptide partitioning to DDX4N1 condensates. In the first model (composition model), the partition free energy for a variant, $v$, is the sum of the individual contributions of each amino acid in the sequence:

$$\Delta G_v^\circ = \sum_j^{20} \beta_j x_j \quad \text{(S13)}$$

where $j$ iterates over the 20 natural amino acids, and $x$ is the number of this type of amino acid in the sequence. The $\beta$-values can therefore be thought of as the energetic contribution ($\Delta G_{part}$) from each type of amino acid, assuming that all these energies are additive.

In order to involve positional information in mutation libraries, a slightly expanded model (substitution model) was used. Here, a $\beta$-value was assigned to each amino acid at all positions in the peptide tile. The equation for the partition energy is, therefore, a double sum with the dummy variable $x_{i,j}$ being 1 when amino acid $j$ is at position $i$, or else it is 0.

$$\Delta G_v^\circ = \sum_i^{L} \sum_j^{20} \beta_{i,j} x_{i,j} \quad \text{(S14)}$$

By subtracting the matrix of the wild type sequence from $x_{i,j}$ and adding a fitting constant to represent the wild type peptide's partition energy, the $\Delta\Delta G = \Delta G_{mutant} - \Delta G_{\mathrm{WT}}$ caused by all amino acid substitutions can be obtained under the assumption that they are all additive. This is similar to Global Multi-Mutant Analysis (GMMA) to identify the effect of amino acid substitutions in folded proteins [5], and we therefore adopt that name:

$$\Delta G_v^\circ = \sum_i^{L} \sum_j^{20} \Delta\Delta G_{i,j} x_{i,j} + \Delta G_{\mathrm{WT}} \quad \text{(S15)}$$

Finally, when no reference wild type sequence was available, the context dependence of amino acids effects were evaluate using a $20 \times 20$ coupling matrix, $M_c$(first,second), where the additional energy related to pairs of amino acids occurring together was fitted.

$$\Delta G_v^\circ = \sum_{j}^{20} \beta_j x_j + M_c(\text{first,second}) \tag{S16}$$

Before any model fitting, the data was split in a ratio of 9:1 for adjusting the hyper-parameters (training set) and testing, respectively. To fit the model, the training set was used to parameterize hyperparameters using 5-fold cross-validation for regularised weighted linear regression in Python. L2 regularisation was used on fitting the 760 $\Delta\Delta G$-values from the mutation library and the $20 \times 20$ coupling matrix. Similar models were used to study the effect of nucleotides in the RNA sequence.

##### Model to relate partitioning into DDX4N1 to intrinsic phase separation propensity

We reasoned that at least a first order correction was needed in order to relate partitioning of peptides into DDX4N1 condensates to how the same peptides would partition to different condensates. Diverse condensates formed *in vitro* behave as hydrophobic solvents of different strength[6] and given condensates formed by completely disordered IDRs, it is unclear to what extent specific interactions should be present[7]. As an example, there is a preference for interactions between the oppositely charged IDRs of HNRNPA1 and FUS compared to homotypic interactions, but this is absent when the charge of HNRNPA1 is changed to that of FUS by mutations[8].

Thus, in the absence of charge, we argued that it would be possible to normalize for the hydrophobicity[1] of the condensate formed by an IDR. We therefore postulate that the partitioning free energy of a peptide $z$ into condensed phase formed by peptides $x$ or $y$ can be described as:

$$\frac{\Delta G(z|x)}{\Delta G(z|y)} = \frac{\Phi_x}{\Phi_y} \tag{S17}$$

where, $\Delta G(x|y)$ is a notation for the partition free energy of peptide $x$ partitioning to a liquid phase of $y$ and $\Phi$ is the hydrophobicity/stickiness of a given phase. This means that peptide will partition with twice the free energy to a condensate that is twice as hydrophobic/sticky.

As proxy for the hydrophobicity of a condensed phase ($\Phi$), we use a measure for the interactions possible within the condensed phase using three simplifying assumptions: The weight to volume concentration of peptides in the condensed phase is roughly constant, all amino acids have the same volume, and interactions between all amino acids are possible. We define the partition free energy density of a peptide $x$ by its partition free energy into a reference peptide $y$, $D(x|y)$, as:

$$D(x|y) \equiv \frac{\Delta G(x|y)}{L_x} \tag{S18}$$

given a large excess of $y$ ($[y] >> [x]$) and $L_x$ is the length (number of amino acid residues) of peptide x. The extrapolation of partitioning of a third peptide $z$ then becomes:

$$\frac{\Delta G(z|x)}{\Delta G(z|y)} = \frac{D(x|y)}{D(y|y)} = \frac{\frac{\Delta G(x|y)}{L_x}}{\frac{\Delta G(y|y)}{L_y}} = \frac{\Delta G(x|y)}{\Delta G(y|y)} \frac{L_y}{L_x} \tag{S19}$$

using equation S17, which is intuitively understood such that a hydrophobic peptide will also form a very hydrophobic condensate, such that it partitions all other peptides more strongly.

By substituting $x$ for $z$ and rearranging the equation above, it then follows that the partition free energy of $x$ to phase separate alone, $\Delta G(x|x)$, can be expressed as (Supplementary Figure 12):

[1]Here we used hydrophobicity for lack of a better term. For phase separation, it has also been called "stickiness". This term comprises all non-electrostatic components of favourable interaction between amino acids.

$$\Delta G(x|x) = \frac{\Delta G(x|y)^2}{\Delta G(y|y)} \frac{L_y}{L_x} \tag{S20}$$

In the context of our experiments, equation S20 shows that the intrinsic phase separation propensity of a client peptide has a square dependence on its measured partition free energy to go into a DDX4N1 condensate:

$$\Delta G(\text{Client}|\text{Client}) = \frac{\Delta G(\text{Client}|\text{DDX4N1})^2}{\Delta G(\text{DDX4N1}|\text{DDX4N1})} \frac{L_{\text{DDX4N1}}}{L_{\text{Client}}} \tag{S21}$$

where, $\Delta G(\text{Client}|\text{DDX4N1})$ can be measured by mRNA-display and $\Delta G(\text{DDX4N1}|\text{DDX4N1})$ is the partition free energy of DDX4N1 obtained by $\Delta G(\text{DDX4N1}|\text{DDX4N1}) = -RTlog\left(\frac{[\text{DDX4N1}]_{\text{condensed}}}{[\text{DDX4N1}]_{\text{dilute}}}\right)$. Similarly, equation S19 allows for extrapolation of how a given peptide partitioning into DDX4N1 would partition to another condensate formed by third peptide, if the partition free energy of the third peptide to DDX4N1 has been measured.

Intuitively, the square dependence can be understood as phase separation originating from two effects: the hydrophobicity of a peptide and the hydrophobicity of the resulting condensate. This is well illustrated for the effect of mutations to a wild type (WT) peptide/protein that does not change the length of the amino acid chain ($L_{\text{Mutant}} = L_{\text{WT}}$). By rearrangement of equation S20, the partition free energy of the mutant into the WT $\Delta G(\text{Mutant}|\text{WT})$ is:

$$\Delta G(\text{Mutant}|\text{WT}) = \sqrt{\Delta G(\text{Mutant}|\text{Mutant})\Delta G(\text{WT}|\text{WT})} \tag{S22}$$

The equation is symmetric such that the $\Delta G(\text{Mutant}|\text{WT}) = \Delta G(\text{WT}|\text{Mutant})$, meaning that a given mutation has equivalent effects on the scaffold and client, making the partioning peptide and the condensate more/less hydrophobic. Therefore, $\Delta G(\text{Mutant}|\text{Mutant}) \propto \Delta G(\text{Mutant}|\text{WT})^2$ which for dilute phase concentration translates to:

$$ln(C_{sat}(\text{Mutant}|\text{Mutant})) \propto \frac{1}{\Delta G(\text{Mutant}|\text{WT})^2} \tag{S23}$$

This is similar to earlier results showing that $ln(c_{\text{sat}}) \propto 1/(n_{\text{Tyr}} n_{\text{Arg}})$, which was rationalised using mean field polymer theory[9]. In our framework, we generalize to interaction between all amino acid by the usage of the energy density of the IDR to extrapolate partitioning in DDX4N1 to other IDR condensate results in the following generalization of proportionality for homotypic phase separation: $ln(c_{sat}) \propto 1/\Delta G^2_{\text{part,DDX4N1}}$.

#### Sequences and primers

pET29a (+) with eMM9-tagged Ddx4N1 CtoA. Cloned between NdeI and XhoI: CATATGGTGCTGGACGTTACCAAGGATCACTGGCTGCCGTACGTGCTGCTGGCGCAGCTGCCGGTGATGGTTCTGTTCCGTAAAGACAACGATGAGGAAGCGAAGAAAGTGGAGTATATTGTTCGTGAGCTGGCGCAAGAATTCGACGGTCTGATCAAGGTTTTTGTGGTTGATATCAACAAAGCGCCGGAAATTGCGAAGAAATACAACATCACCACCACCCCGACCGTGGCGTTCTTTAAGAACGGCGAGCTGAAAAGCGTTTTTACCGGCGCGATTAGCAAGGACCAGCTGCGTGATGAAATCCTGAAATACCTGGGTCATCATCATCATCACCACGGTAGCGGCAGCGGCAGCGGTAGCGGTAGCGAGAACCTGTATTTCCAGGGCATGGGTGACGAAGATTGGGAGGCGGAAATTAACCCGCACATGAGCAGCTATGTTCCGATCTTTGAGAAAGATCGTTACAGCGGTGAAAACGGCGACAACTTCAACCGTACCCCGGCGAGCAGCAGCGAGATGGACGATGGCCCGAGCCGTCGTGACCACTTCATGAAGAGCGGTTTTGCGAGCGGCCGTAACTTTGGTAACCGTGATGCGGGTGAAGCGAACAAACGTGACAACACCAGCACGATGGGTGGTTTCGGCGTTGGCAAGAGCTTCGGTAACCGTGGCTTTAGCAACAGCCGTTTCGAAGACGGCGATAGCAGCGGTTTTTGGCGTGAGAGCAGCAACGATGCGGAAGACAACCCGACCCGTAACCGTGGTTTCAGCAAACGTGGCGGTTATCGTGATGGCAACAACAGCGAGGCGAGCGGTCCGTACCGTCGTGGCGGTCGTGGCAGCTTTCGTGGTGCGCGTGGCGGTTTCGGTCTGGGTAGCCCGAACAA

CGACCTGGATCCGGACGAAGCGATGCAACGTACCGGCGGTCTGTTTGGCAGCCGTCGTCCGGTGCTGAGCGGTACCGGTAACGGTGACACCAGCCAAAGCCGTAGCGGTAGCGGTAGCGAGCGTGGCGGTTACAAGGGTCTGAACGAGGAAGTTATCACCGGTAGCGGCAAAAACAGCTGGAAGAGCGAGGCGGAAGGCGGTGAAAGCTAACTCGAG

`Ddx4N1 CtoA`: GMGDEDWEAEINPHMSSYVPIFEKDRYSGENGDNFNRTPASSSEMDDGPSRRDHFMKSGFASGRNFGNRDAGEANKRDNTSTMGGFGVGKSFGNRGFSNSRFEDGDSSGFWRESSNDAEDNPTRNRGFSKRGGYRDGNNSEASGPYRRGGRGSFRGARGGFGLGSPNNDLDPDEAMQRTGGLFGSRPVLSGTGNGDTSQSRSGSGSERGGYKGLNEEVITGSGKNSWKSEAEGGES

`Pep6_lib_fw` (mutated amino acids in lower case): TTAACTTTAAGAAGGAGATATACATATGcgtttcgaagacggcgatagcagcggtttttggcgtgagagcagcaacgatgcggaagacaa

`Pep6_lib_rev` (mutated amino acids in lower case): GGATAGCTACCGCTACCgctgttgttgccatcacgataaccgccacgtttgctgaaaccacggttacgggtcgggttgtcttccgcatcg

`GS3an_R36`: TTTCCGCCCCCCGTCCTAGCTGCCGCTGCCGCTGCC

`Fivep_flank`: TAATACGACTCACTATAGGGTTAACTTTAAGAAGGAGATATACATATG

`Threep_flank_HA`: TTTCCGCCCCCCGTCCTAGCTGCCGCTGCCGCTGCCTGCATAATCCGGAACATCATACGGATAGCTACCGCTACC

`Rd1T7g10M_F71` + phasing: ACACTCTTTCCCTACACGACGCTCTTCCGATCT(N)_(0-8)TAATACGACTCACTATAGGGTTAACTTTAAGAAGGAGA

`an13Rd2_R49` + phasing: GACTGGAGTTCAGACGTGTGCTCTTCCGATCT(N)_(0-8)TTTCCGCCCCCCGTCCT

`Rd2N(701)P7_R52` + multiplexing barcode: CAAGCAGAAGACGGCATACGAGAT(TCGCCTTA)GTGACTGGAGTTCAGACGTG

`P5S(502)Rd1_F57` + multiplexing barcode: AATGATACGGCGACCACCGAGATCTACAC(CTCTCTAT)ACACTCTTTCCCTACACGAC

| Sample | Total reads | Merged reads | Mapped reads |
|---|---|---|---|
| Display LLPS 140 | 1,068,640 | 1,068,640 | 645,269 |
| mRNA LLPS 140 | 1,295,529 | 1,295,529 | 429,836 |
| Display control 140 | 1,223,434 | 1,223,434 | 823,360 |
| mRNA control 140 | 801,776 | 801,776 | 384,089 |
| Display LLPS 158 | 1,187,057 | 1,187,057 | 776,302 |
| mRNA LLPS 158 | 1,191,221 | 1,191,221 | 490,263 |
| Display control 158 | 1,103,307 | 1,103,307 | 734,922 |
| mRNA control 158 | 650,312 | 650,312 | 306,822 |

**Table S1:** Sequencing statistics for the tiling library.

| Sample | Total reads | Merged reads | Reads of correct length |
|---|---|---|---|
| Display LLPS | 156,506,905 | 67,322,896 | 63,548,990 |
| mRNA LLPS | 186,713,009 | 59,118,197 | 49,947,771 |
| Display control | 155,008,791 | 93,984,207 | 90,360,925 |
| mRNA control | 100,190,128 | 42,067,683 | 36,114,813 |

**Table S2:** Sequencing statistics for the first measurement of the mutation library.

| Sample | Total reads | Merged reads | Reads of correct length |
|---|---|---|---|
| Control Display 1 | 19,294,036 | 11,988,970 | 10,888,299 |
| Control Display 2 | 23,540,235 | 14,698,024 | 13,341,503 |
| Control mRNA 1 | 20,687,495 | 8,665731 | 6,624,173 |
| Control mRNA 2 | 37,498,756 | 15,236,179 | 11,612,596 |
| Control pur 1 | 60,321,005 | 21,756,967 | 15,354,958 |
| Control pur 2 | 70,047,841 | 25,103,674 | 17,796,570 |
| LLPS Display 1 | 71,978,308 | 17,510,300 | 15,301,835 |
| LLPS Display 2 | 72,584,805 | 27,301,027 | 24,043,644 |
| LLPS mRNA 1 | 88,135,303 | 25,809,001 | 19,572,304 |
| LLPS mRNA 2 | 89,380,343 | 26,517,893 | 19,906,843 |
| LLPS pur 1 | 50,747,381 | 13,174,457 | 8,990,215 |
| LLPS pur 2 | 67,865,585 | 17,359,945 | 11,848,672 |

**Table S3:** Sequencing statistics for the second measurement of the mutation library.

| Sample | Total reads | Merged reads | Mapped reads |
|---|---|---|---|
| Control Display 1 | 123,863,370 | 117,840,378 | 104,383,295 |
| Control Display 2 | 92,555,662 | 88,063,342 | 78,138,850 |
| Control mRNA 1 | 127,547,518 | 110,745,138 | 97,513,571 |
| Control mRNA 2 | 152,108,027 | 128,912,292 | 112,166,493 |
| Control pur 1 | 194,943,764 | 160,513,402 | 136,915,411 |
| Control pur 2 | 143,121,647 | 120,065,735 | 103,062,120 |
| LLPS Display 1 | 113,578,830 | 105,580,653 | 92,731,531 |
| LLPS Display 2 | 122,587,015 | 114,088,323 | 100,836,577 |
| LLPS mRNA 1 | 133,909,387 | 113,682,039 | 100,002,766 |
| LLPS mRNA 2 | 118,890,716 | 100,857,535 | 89,182,316 |
| LLPS pur 1 | 123,792,492 | 104,434,822 | 90,036,388 |
| LLPS pur 2 | 111,795,820 | 94,158,151 | 81,109,900 |

**Table S4:** Sequencing statistics for the DisProt library.

| Library | Data formatting | Model | Regularisation | $R^2$ training | $R^2$ test | $R^2$ total |
|---|---|---|---|---|---|---|
| Pep6Lib | Composition | OLS | N/A | N/A | N/A | 0.180 |
| Pep6Lib | One-hot (substitutions) | Ridge (L2) | $\lambda = 2 \cdot 10^3$ | 0.283 | 0.249 | 0.284 |
| DPLib | Composition (single and double) | Ridge (L2) on doubles | $\lambda = 2 \cdot 10^5$ | 0.428 | 0.428 | 0.429 |

**Table S5:** Performance of models.

| Gene | Reference |
|---|---|
| AFF4 | Guo et al., 2020 [10] |
| AR | Zhang et al., 2023 [11] |
| DDX3X | Shen et al., 2022 [12] |
| EIF4B | Swain et al., 2024 [13] |
| EWSR1 | Johnson et al., 2024 [14] |
| ESR1 | Nair et al., 2019 [15] |
| FMR1 | Kim et al., 2019 [16] |
| FUS | Patel et al., 2015 [17] |
| HNRNPA1 | Molliex et al., 2015 [18] |
| HNRNPA2B1 | Ryan et al., 2018 [19] |
| MAPT | Wegmann et al., 2018 [20] |
| MBP | Aggarwal et al., 2013 [21] |
| POLR2A | Boehning et al., 2018 [22] |
| TARDBP | Molliex et al., 2015 [18] |

**Table S6:** References for human genes encoding proteins containing IDRs that are highly partitioned to DDX4N1 condensate according to our CPmD data and have previously been described to form condensate in purified form.

| Gene | Reference |
|---|---|
| BRCA1 | Qin et al., 2023 [23] |
| BRCA2 | Skobelkina et al., 2024 [24] |
| GLI3 | Han et al., 2025 [25] |
| LAT | McAffee et al., 2022 [26] |
| NOLC1 | Courchaine et al., 2022 [27] |
| PGR | Muñoz-Gil et al., 2020 [28] |
| PPARGC1A | Pérez-Schindler et al., 2021 [29] |
| PRC1 | Eeftens et al., 2021 [30] |
| SNRNP70 | Hu et al., 2022 [31] |
| SRRM1 | Niedner-Boblenz et al., 2024 [32] |
| TOB1 | Perez et al., 2024 [33] |

**Table S7:** References for genes encoding proteins containing IDRs that are highly partitioned to DDX4N1 condensate according to our CPmD data, and have previously been described to participate in formation of biomolecular condensates.

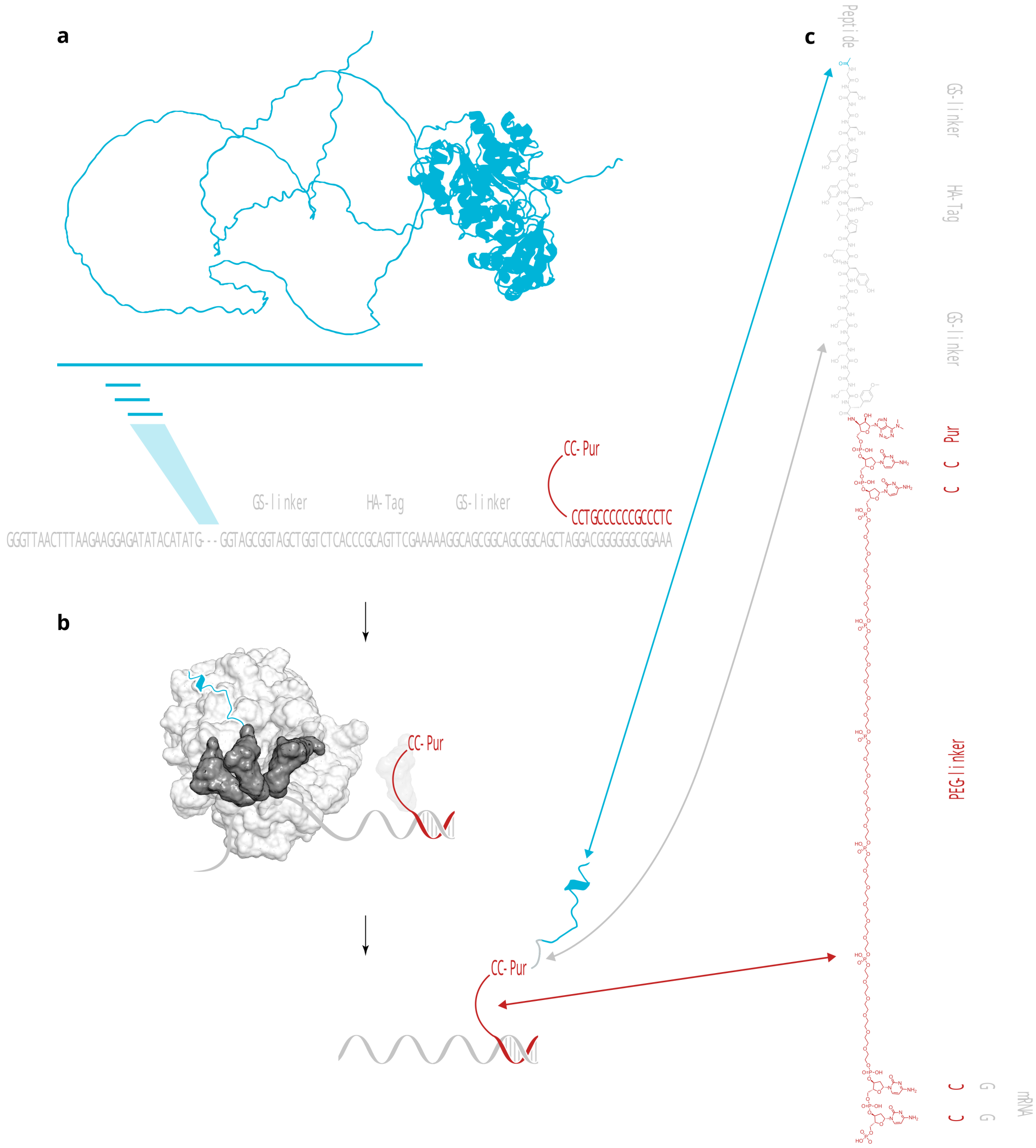


**Figure S1: Overview of the mRNA-display construct used in CPmD. a,** DDX4 (in blue) was tiled into peptides that were encoded in synthetic genes containing all elements needed for transcription. Glycine/serine linkers are also encoded together with an HA-tag for purification. A puromycin containing linker (red) is annealed to the transcribed mRNA and covalently attached via ligation at the 5' end (covalent product not shown). **b,** The resulting fusion mRNA-linker is used for translation using the PURE-system[34] where release factors 1, 2, and 3 are omitted. The ribosome then stalls at the stop codon which allows the puromycin to react with the nascent peptide to form the peptide-linker-mRNA fusion shown in **c**.

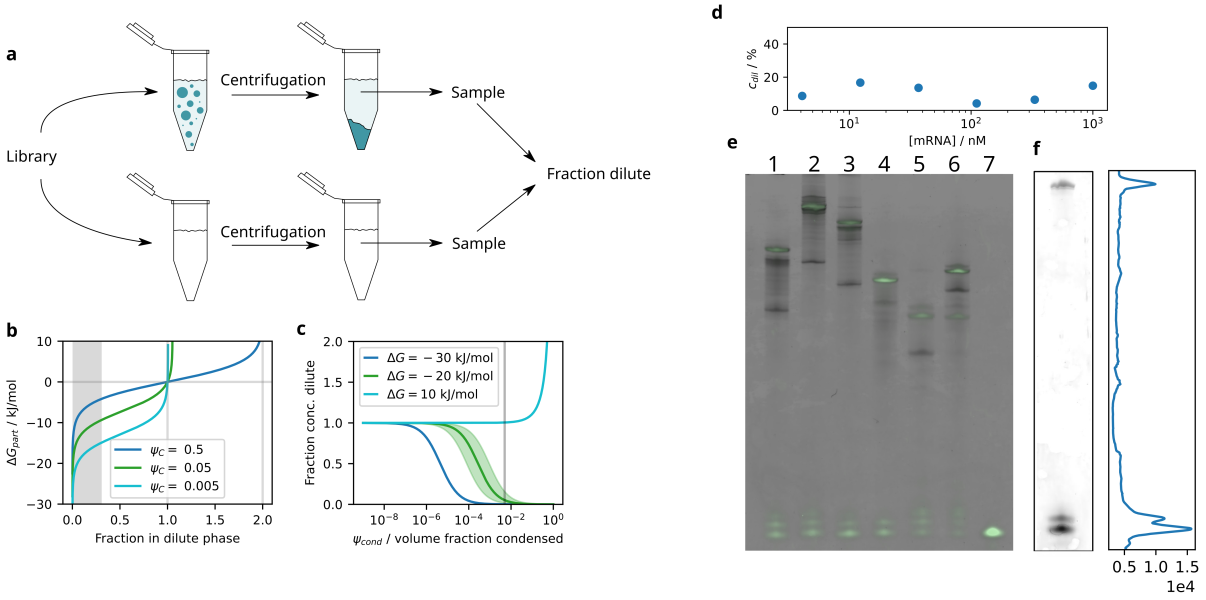


**Figure S2: CPmD can measure partition free energies of both peptides and full proteins.** **a,** Two parallel experiments are performed where either the mRNA-display library dissolved in low salt buffer is added to a concentrated protein stock to induce phase separation or a buffer control without protein. Centrifugation is then used to sediment the condensed phase, and a sample from the dilute phase is reverse transcribed. qPCR estimates of cDNA concentration in the samples together with sequencing reads are used to assign a concentration to all library members. The ratio of the two concentrations is the fraction remaining in the dilute phase. **b**, Using this value, the partition free energy is calculated using mass action, by assuming a condensed phase volume fraction, $\Psi_C$ (typically of the order of 0.005). **c**, $\Psi_C$ was chosen such that only a small fraction of the library was remaining, which made analysis simpler (See equation S8). **d**, The fraction of the concentration remaining in the dilute phase of a control mRNA molecule was roughly constant across two orders of magnitude of total concentration, showing that the interaction is indeed partitioning and not specific binding. **e**, CPmD is also applicable to entire proteins and domains: sample 1) [160-199] of Ddx4N1, 2) $\alpha$-synuclein, 3) [30-110] $\alpha$-synuclein, 4) HiBit, 5) Single StrepII, 6) Twin StrepII, 7) puromycin linker alone. **f** A critical step in the mRNA-display protocol is the attachment of the puromycin linker, which is also possible for the full DDX4N1 domain. This construct was also successfully translated (data not shown).

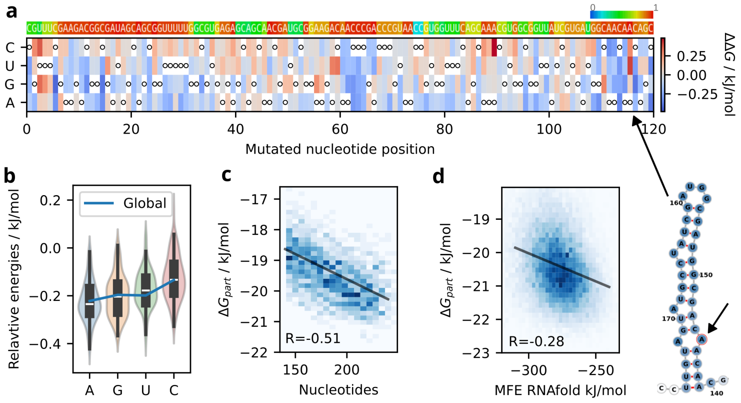


**Figure S3: Long, less structured, and purine-rich mRNA partitions stronger.** **a**, A GMMA-model fit of mRNA partitioning from the mutation library shows effects of mutations in the sequence. On top, the base pairing probability from RNAfold[35]. A strong effect disfavoring partitioning is seen from A145U, which corresponds to a predicted mismatch in a hair-pin structure, showing how secondary structure formation excludes mRNA from the condensate. **b**, The average effect on partitioning of each type of nucleotide in a similar fit as in a, which includes WT nucleotides rather than relative $\Delta\Delta G$. **c**, The length of the mRNA also has a strong effect on partitioning, which is shown here for the tiling library of DDX4N1 with various peptide lengths. **d**, Predicted minimum free energy (MFE) of all mRNAs used for fit in panel a shows that stronger folding of the mRNA disfavours partitioning - an effect seen for all libraries.

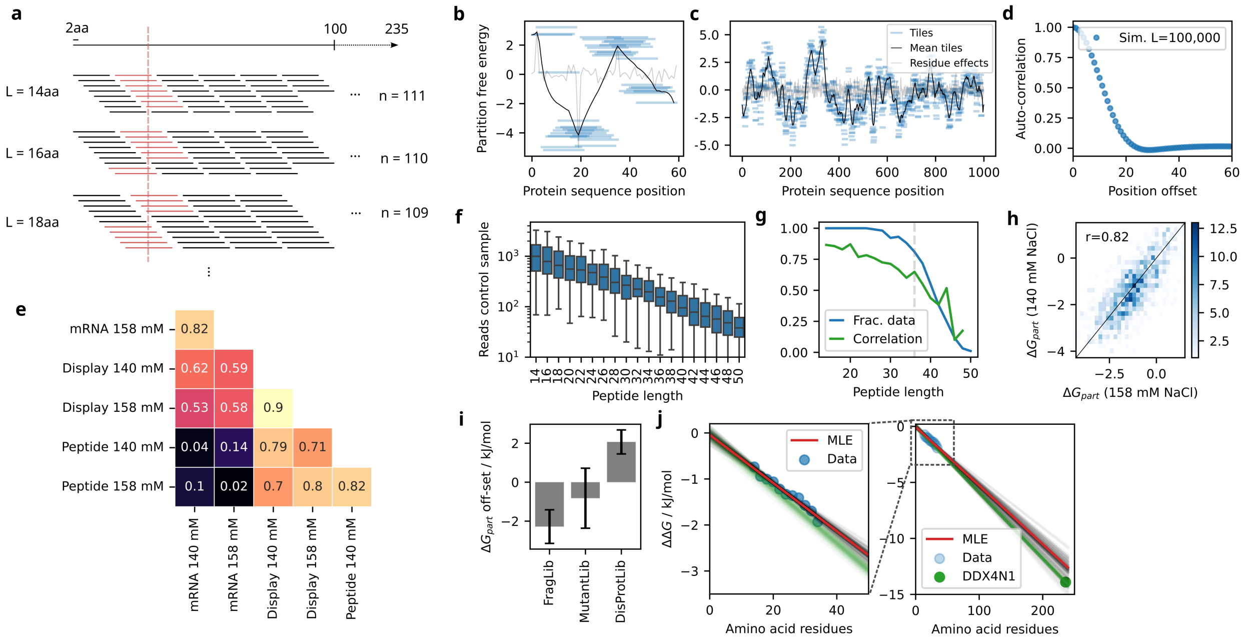


**Figure S4: Tiling library of DDX4N1.** **a**, Design of tiling library and principle of computing a running mean to interpret the data. Tiles of 14 to 50 amino acids of even lengths were constructed to start from every other position in DDX4N1. The running mean was then calculated for every position of DDX4N1 by computing the mean partition free energy of all peptides containing the amino acid at that position. **b**, A simulated example of such a running mean calculation. In grey, the contribution from amino acids at all positions are plotted. Tile energies obtained by adding the amino acid energies together are shown in blue. The running mean is shown in black, and is much smoother than the underlying causes for partitioning. **c**, Simulation of the example in b, but with random amino acid contributions to the partition free energy. Even in such a case, the running mean is observed to have pronounced features of hills and valleys. **d**, A simulation of a hypothetical protein of 100,000 residues shows that the auto-correlation in the running mean is very close to the peptide tile length used, 16 amino acids. **e**, Pearson's correlation coefficient between two CPmD replicates that were performed at slightly different NaCl concentration causing different (but similar) dense phase volume fractions done for the fragment library for two samples with mRNA and mRNA-display. The difference between the two, "Peptide", is also shown. This shows that the obtained partition free energies correlate with the mRNA-display samples, but not with the mRNA itself. This supports our view that we can disentagle mRNA and peptide effects. **f**, The shorter constructs in the library took up many more sequencing reads than the longer. **g**, The bias towards shorter sequences highlighte din panel f caused a declining quality of the data for the longer peptide fragments. **h**, Correlation shown for the two 14 amino acid peptide replicates. **i**, The absolute scale of the resulting partition free energies is not well defined, because it relies on the qPCR quantification of total library concentration. To correct for this, a global composition model fit of all libraries measured was performed. The fit resulted in different absolute off-set for the three libraries. For the tiling library of DDX4N1 (FragLib) and multi-mutant library of the 40 amino acid tile of DDX4N1 (MutantLib), the peptide energies were only corrected using pure mRNA as the reference such that the linker was not accounted for, as it was done in the DisProt database library (DisProtLib). The linker partitioning is therefore unfavorable. This could be due to the formation of a heteroduplex between the linker and RNA that might be unfavorable, given that single stranded nucleic acids have been shown to partition preferentially[36]. The two first libraries therefore likely have the same intercept, with the value for the FragLib being the better estimate. The off-set used for the MutantLib in subsequent analysis is the weighted mean between the FragLib and MutantLib values. The off-set for the DisProLib likely originates from the GS-linkers and HA-tag, which are present on all peptides in our protocol. The estimated off-set is within error of the partition free energy for 2xGS-HA-3xGS predicted by our composition model ($-1.13$ kJ/mol, p=0.15). **j**, The off-set estimate mean partition free energy of the peptides from the FragLib ($-2.28 \pm 0.86$ kJ/mol) was used as a prior for the intercept in linear regression of the length dependence of the peptide partitioning. Data is shown with a pre-correction for $-2.28$ kJ/mol.

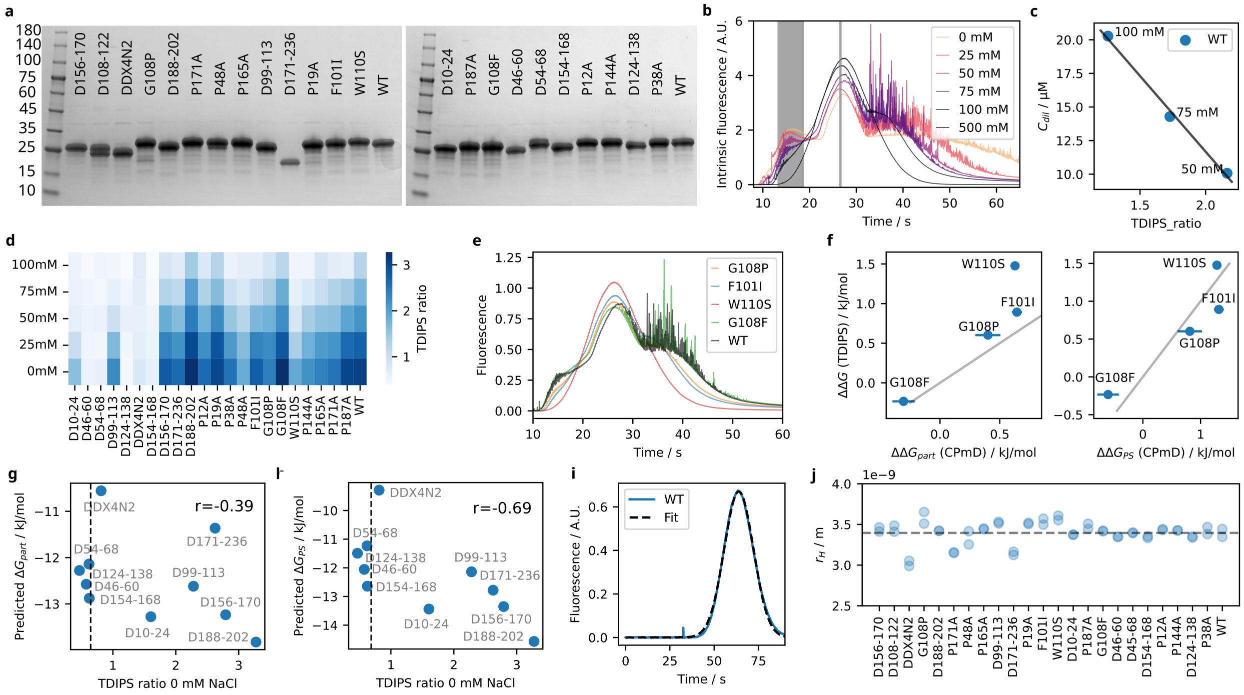

**Figure S5: Measurements of condensate formation of purified DDX4N1 mutants using TDIPS.** **a**, SDS-PAGE gel of purified mutants of DDX4N1. The deletion mutant D108-122 showed a double band and was omitted from further analysis. **b**, Taylor-Dispersion Induced Phase Separation (TDIPS)[37] of DDX4N1 at different concentrations of NaCl in the surrounding buffer. The degree of phase separation was quantified by the ratio between the integrated area shown in grey between 10 and 20 seconds representing dissolved condensates transported in the laminar flow, and the height of the monomer peak at ∼26 seconds. **c**, The known dilute phase concentration of DDX4N1 was used to calibrate a standard curve in order to relate TDIPS ratio to dilute phase concentration. **d**, Measured TDIPS ratios for all purified variants at different concentrations of NaCl. Higher ratio (more blue) means lower dilute phase concentration and stronger phase separation. **e**, TDIPS curves for four point mutants of DDX4N1. **f**, Using the standard curve in panel c, the TDIPS ratios of the mutants were converted to changes in phase separation free energy $\Delta\Delta G$. These values were then compared with the GMMA-fitted values from the mutant library $\Delta\Delta G_{part}$ (left plot). The correlation improved when correcting these values using the partitioning framework described above, such that the values instead should report on phase separation rather than partitioning $\Delta\Delta G_{PS}$ (right plot). **g**, The predicted partition free energies of the deletion mutants correlated with the measured TDIPS ratios. Vertical interrupted line indicates the bottom of the dynamic range in the TDIPS experiments. **h**, Predicting instead the phase separation free energy, the correlation improves supporting the view that the deletions also affect the "hydrophobicity" of the resulting condensates. **i**, Flow Induced Dispersion Analysis (FIDA)[38] was also performed to measure the hydrodynamic radius ($r_H$) of the mutants. **j**, Hydrodynamic radius of all purified mutants.

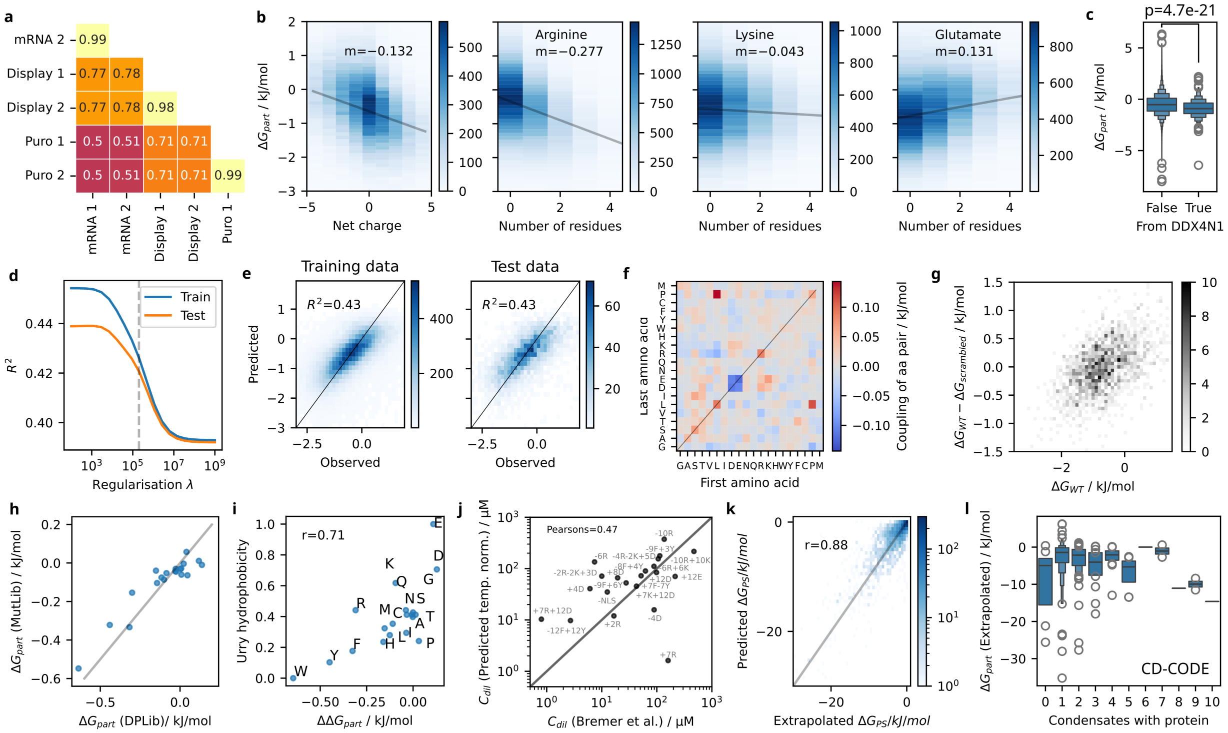

**Figure S6: DisProt library and fitting of the composition model.** **a**, Correlations between the measured partition free energies of mRNA, mRNA-puromycin linker (puro), and mRNA-display (Display) for two replicates. **b**, Correlation between peptide partition free energy and Net Charge Per Residue (NCPR) and three of the charged amino acids. **c**, Partition free energies of all peptides in the DisProt library depending on if the are from DDX4N1 or not. **d**, Fitting of the composition model via L2 regularisation. The regularisation was only applied to pair terms for neighbouring residues (panel f) and not single amino acids. **e**, Performance of the model on training and test data. **f**, Matrix for pair terms for neighboring amino acids. **g**, For peptides where scrambled versions were included, the only parameter correlating with the difference between WT and Scrambled partition free energy was the energy of the WT. **h**, The fitted single amino acid values for the composition model fitted here to the DisProt library correlated well with the values obtained from the DDX4N1 mutant library. **i**, Correlation between single amino acid energies from the composition model and the Urry hydrophobicity scale[39]. **j**, Performance of the composition model together with the partitioning framework (for 20°C) in predicting data on hnRNPA1 mutants measure at 4°C from Bremer et al. 2022[40]. All predicted dilute phases are scaled based on the ratio of measured dilute phase concentrations of hnRNPA1 WT at 4 and 20°C. **k**, Agreement between extrapolated partition free energy of the IDRs in the DisProt data based the average partitioning of the tiles and the partition free energy predicted directly from the composition model. **l**, Correlation between the extrapolated partition free energy from the DisProt library and the number of condensates reported to contain them in the CD-CODE database[41].

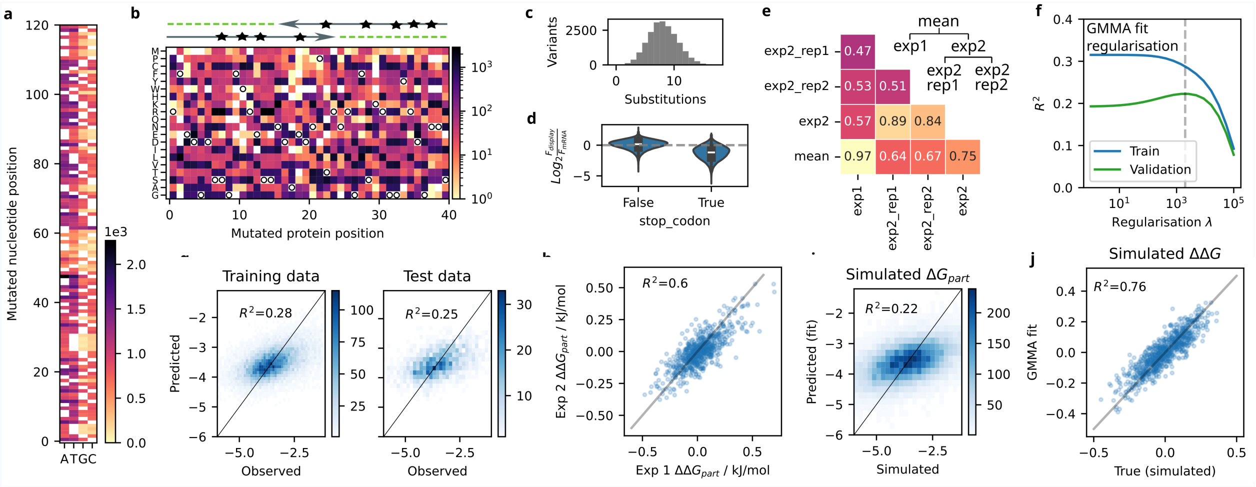


**Figure S7: Multi-mutant library of DDX4N1 100-140 and GMMA fitting.** **a**, We created a mutation library of a 40 amino acid long peptide tile of DDX4N1 (residues 100-139) from two partially overlapping doped oligonucleotides. The number of mutated DNA bases at each position in the mutated region is shown. The overlap required from assembly of the oligonucleotides is visible at position ∼50-60, were there is a lower frequency of mutations due to the requirement for base pairing at 37°C. **b**, The resulting amino acid substitutions are shown, with WT amino acids indicated by circles. **c**, The distribution of the number of amino acid substitutions in the variants of the library. **d**, The frequency of reads after translation of the mRNA and subsequent affinity purification via the HA-tag on the peptide for variants with or without a premature stop codon. Stop codons cause a significant decrease in abundance, but the difference is only about two-fold. This suggests that stop codons are frequently read through by the ribosomes. **e**, Correlation between the different replicates of peptide partition free energies in the library. **f**, GMMA fitting was done as described above in the section on Linear methods. Here, training and validation coefficient of determination, $R^2$, is shown while tuning L2 regularisation to control the fitted $\Delta\Delta G$-values to explain the partitioning of the multi-mutants. **g**, Resulting correlation between training and test splits of the data. **h**, Correlation between $\Delta\Delta G$-values obtained from fitting on the two main replicates (exp1 and exp2). **i**, The inability of the GMMA-fit to capture the amplitude of the data (i.e. "flatness" in panel g) causes a low $R^2$-value while the correlation is much better. Simulated peptide partitioning data with high amounts of error on individual partition free energy measurements show the same "flatness" of the fit. **j**, However, the resulting fitted $\Delta\Delta G$-values accurately correlate with the true values used for the simulations. The correlation coefficient matches the one obtained between the two replicates (exp1 and exp2) in panel h, showing that the "flatness" in our GMMA fit on the CPmD data is caused by noisy measurements. However, the obtained $\Delta\Delta G$-values are likely accurate.

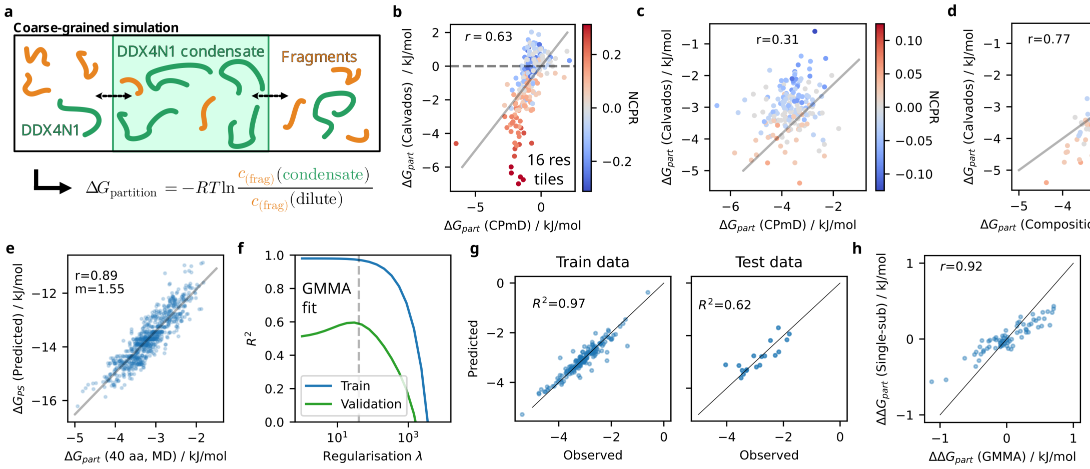


**Figure S8: Coarse-grained molecular dynamics simulations of partitioning experiments.** **a**, Using the CALVADOS force-field for simulations of intrinsically disordered proteins, we simulated an experiment resembling CPmD. An excess of DDX4N1 molecules were simulated in a slab-geometry and formed a condensate, and the partitioning of several peptide fragments/tiles was followed. The partition free energies of the peptides were calculated from the average concentration in the dilute and condensed phase across the simulation time. **b**, Comparison between CPmD data and CALVADOS simulations for 16 amino acids long tiles of DDX4N1. **c**, Almost 200 multi-mutant peptides from the multi-mutant library were simulated with CALVADOS, and the resulting correlation is shown. The CPmD data from this library contained significant noise (error bars not shown for clarity), which made comparison difficult but still a good correlation was observed. GMMA extracted single amino acid effects correlated better (Fig. 3 in main manuscript). **d**, Across the simulations, the charge of the peptides strongly correlated with partitioning, which was less the case for CPmD data. Using the fitted composition model on CPmD data, we found good agreement with the CALVADOS partitioning of the multi-mutants. This showed that the net charge of the peptides likely significantly influence partitioning in the simulations, as it is not considered in the composition model. **e**, We also simulated single amino acid substitution variants of the 40 amino acid peptide (Fig. 3, main manuscript). A neural-network model trained to predict intrinsic phase separation based on CALVADOS simulations[42] correlated well with the simulated partition free energies. This supported our hypothesis that partitioning to DDX4N1 condensates and intrinsic phase separation propensity is strongly linked. The slope between the two values is not unity, as would also be expected based on our derived partitioning framework of phase separation. **f**, Using the set of simulated data of both multi-mutant and single substitution partition free energies, we performed GMMA analysis to extract the single substitution effects from the multi mutant data. Regularisation on the $\Delta\Delta G$-values is shown. **g**, Correlation between training and test splits for the chosen regularisation strength. **h**, Correlation between the simulated effects of single amino acid substitutions and those fitted using GMMA. This computational GMMA experiment demonstrates that it is possible to extract single variant effects from the global analysis of the multi mutant library and supports the model of fitted additive effetcs.

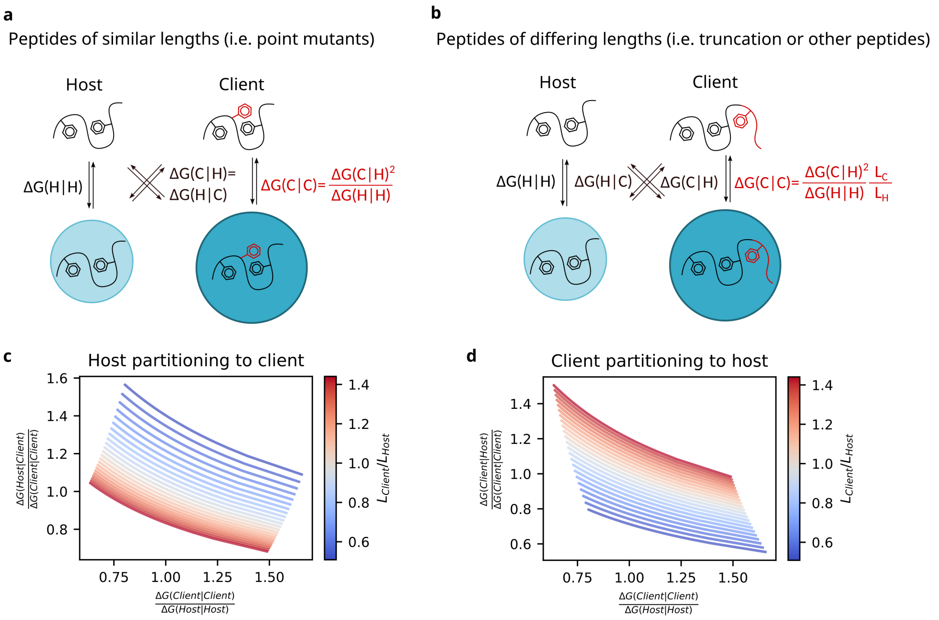


**Figure S9: Partitioning framework for phase separation.** **a**, We hypothesize that phase separation of peptides can be split into two contributions: 1) the inherent driving force (i.e. "hydrophobicity" or "stickiness") for a small amount of peptide to partition to a condensate formed by some other components, which are in large excess. 2) the "hydrophobicity" of that resulting peptide rich phase that affects how other peptides are partitioned. A simple assumption is that the "hydrophobicty" of a condensate is directly proportional to the energy density of the peptide forming it (defined as the inherent energy of the peptide to partition to some reference condensate, $\Delta G$, divided by the number of amino acids in the peptide, $L$). For point mutants, where the length of the peptide stays the same, this means that the free energy of partitioning is the same for either the client peptide partitioning into a large excess of the host or the host partitioning into a large excess of the client. These assumptions then results in a square dependence of the intrinsic phase separation free energy of the client peptide on its inherent "hydrophobcity" (see Supplementary Methods above). **b**, For the more general case with peptides of different lengths, there is an asymmetry between the free energy of the client partitioning in an excess of the host and the reverse situation. Intuitively, a very long client peptide might have many residues driving strong partitioning to a condensate formed by a host peptide. However, if it also contains many unfavorable amino acids, a condensate formed by that client protein will not be very "hydrohobic" and therefore not be able to recruit the host peptide as strongly. Such asymmetry is shown for a client partitioning to the host, **c**, and the host partitioning to the client, **d**, as a function of relative free energies for phase separation and length difference.
